# Local neuronal secretory trafficking dynamics revealed with zapERtrap: a light-inducible ER release system

**DOI:** 10.1101/2020.09.02.278770

**Authors:** Ashley M. Bourke, Samantha L. Schwartz, Aaron B. Bowen, Amos Gutnick, Thomas L. Schwarz, Matthew J. Kennedy

## Abstract

For normal synapse and circuit function, neurons must regulate the abundance and localization of transmembrane receptor, channel and adhesion proteins over vast cellular expanses, including remote sites in dendrites and axons. Whether the secretory network can support long-range trafficking of synaptic proteins synthesized in the cell body or precise trafficking of locally generated proteins at remote sites remains poorly characterized. We developed an approach for locally triggering secretory trafficking from specific subcellular domains to explore the rate, activity dependence and cargo-specificity of central and remote trafficking networks. Surprisingly, different postsynaptic proteins processed in the cell body were transported deep into dendrites, but with strikingly different kinetics, spatial distributions and activity dependencies. Proteins locally processed in dendrites were broadly dispersed prior to surface insertion, but could be directed locally to synapses. These results provide a novel interrogation of compartmentalized trafficking and reveal basic principles for protein targeting in complex cellular environments.

## Introduction

Integral membrane and secreted proteins are synthesized, processed and delivered to the appropriate subcellular location through a complex set of cellular organelles collectively termed the secretory network. In eukaryotes, the early secretory network is principally defined by the endoplasmic reticulum (ER), the ER-Golgi intermediate compartment (ERGIC) and the Golgi apparatus (GA). In most cells, the physical arrangement of these organelles favors centripetal movement of proteins toward perinuclear GA after they leave the ER. Following GA processing, proteins are sorted and distributed to their appropriate subcellular addresses in distinct classes of mobile vesicle carriers (for reviews see Barlowe and Miller, 2013; Lee et al., 2004; Lippincott-Schwartz et al., 2000).

Large and architecturally complex cells such as neurons face the unique challenge of regulating the abundance and localization of diverse membrane proteins and secreted factors throughout elaborate cellular processes that can project long distances, in some cases several centimeters or meters from the cell body (Bourke et al., 2018; Kennedy and Hanus, 2019; Ramírez and Couve, 2011). How protein demand is satisfied at remote sites in large, compartmentalized cells such as neurons remains a fundamental question in cell biology. In the neuronal cell body, or soma, the general architecture and organization of the secretory organelles resembles conventional cells where secretory trafficking from the somatic ER to the perinuclear GA occurs (Horton and Ehlers, 2003; Horton et al., 2005; Torre and Steward, 1996). However, whether subsequent, post-GA trafficking is restricted to the soma or can be efficiently directed over long distances into dendrites via active vesicular transport mechanisms remains unclear. While direct experimental data is lacking, computational modeling suggests that long-range delivery of somatically processed cargoes to dendrites may be too slow, taking many hours to days, to efficiently accommodate protein demand in remote regions (Williams et al., 2016). Indeed, evidence is accumulating for a distinct, local secretory processing network that could satisfy protein demand in neuronal dendrites. For example, ER-bound ribosomes and mRNAs encoding integral membrane and secreted proteins have been detected in dendrites, implying ongoing local translation of proteins at the dendritic ER (Cajigas et al., 2012; Cui-Wang et al., 2012; Wu et al., 2017). Experiments directly visualizing ER exit and accumulation in the ERGIC established that early trafficking events can occur in dendrites (Bowen et al., 2017; Hanus et al., 2014; Horton and Ehlers, 2003). However, whether subsequent protein trafficking to the cell surface is similarly restricted to a specific dendritic region as cargoes progress through sparsely distributed and highly mobile dendritic organelle networks has remained largely overlooked (Bowen et al., 2017; Cui-Wang et al., 2012). The extent to which the somatic secretory network could also support dendritic protein delivery through direct, long-range vesicular transport remains similarly unclear. Some evidence suggests that surface delivery of postsynaptic receptor proteins is largely restricted to the cell body, with subsequent dendritic localization achieved through lateral diffusion (Adesnik et al., 2005).

To address these issues, new tools allowing precise control over where and when proteins begin their journey through the secretory network are required. Here we introduce a new approach, zapalog-mediated ER trap (zapERtrap), to investigate secretory processing and trafficking from user-defined subcellular domains. ZapERtrap leverages a previously described photosensitive protein dimerizer molecule to sequester protein cargoes within the ER and trigger their release and forward trafficking with a brief (millisecond) pulse of near UV light (Gutnick et al., 2019). Because light can be precisely steered over user-defined cell populations or even subcellular domains, protein trafficking can be initiated with exceptional spatial and temporal control. Combined with sensitive detection methods to monitor surface delivery in real time, we characterized where and when two archetypal synaptic proteins, the cell adhesion molecule neuroligin1 (NL1) and GluA1 AMPA receptors, appear at the cell surface when released from the ER in different subcellular domains. We demonstrate NL1 and GluA1 locally released from the somatic ER are subsequently inserted into the somatic plasma membrane but are also transported deep into the dendritic arbor before surface presentation. For NL1, but not GluA1, the timing, but not total amount of surface delivery was strongly influenced by neural activity. Conversely, activity strongly influenced the total amount of GluA1 delivered to the cell surface but had little effect on the timing. Locally releasing these cargoes from dendritic ER resulted in surprisingly diffuse overall surface delivery, with no enrichment of total surface protein near the site of ER release. While total surface delivery was widespread, we observed an enrichment of cargo at synaptic compartments near sites of ER release. Finally, select cargoes were prevented from entering axons through robust and previously unappreciated surface insertion at the axon initial segment (AIS). Together these data provide the first characterization of the spatiotemporal dynamics of secretory trafficking from distinct subcellular domains. More broadly, zapERtrap opens the door to previously unapproachable questions concerning how proteins are processed, trafficked and secreted in space and time in complex cellular environments.

## Results

### Developing and validating an approach for light-triggered protein secretion

We sought to engineer a system that would allow us to conditionally sequester diverse cargo molecules in the ER, precisely trigger their release at user-defined subcellular regions and visualize their post-ER trafficking itineraries as they progress through the secretory pathway to the plasma membrane. Our system, which we dubbed “zapalog-mediated ER trap” or “zapERtrap”, relies on a small-molecule protein dimerizer “zapalog”. Zapalog consists of the antibiotic trimethoprim (TMP) tethered to a synthetic ligand of FK506-binding protein (SLF) through a photocleavable linker (Gutnick et al, 2019). Thus, zapalog dimerizes FK506-binding protein (FKBP), which binds to SLF, and dihydrofolate reductase (DHFR), which binds to TMP. Brief (millisecond) illumination with low intensity 405 nm light disrupts the photolabile linker bridging SLF and TMP to rapidly dissociate FKBP and DHFR. To design a system for light-inducible ER-release, we fused DHFR to the lumenal domain of several different integral membrane secretory cargo molecules (Fig. 1A). To sequester these molecules in the ER, we targeted FKBP-XFP (where XFP is either EGFP or mCh) to the ER and appended a C-terminal “KDEL” ER retention motif. Thus, in the presence of zapalog, FKBP-XFP-KDEL dimerizes with DHFR-fused cargo molecules, trapping them in the ER. Photocleaving zapalog with 405 nm light rapidly liberates the cargo molecule from the ER retention module (FKBP-XFP-KDEL), allowing forward trafficking to proceed (Fig. 1A).

**Figure 1.**
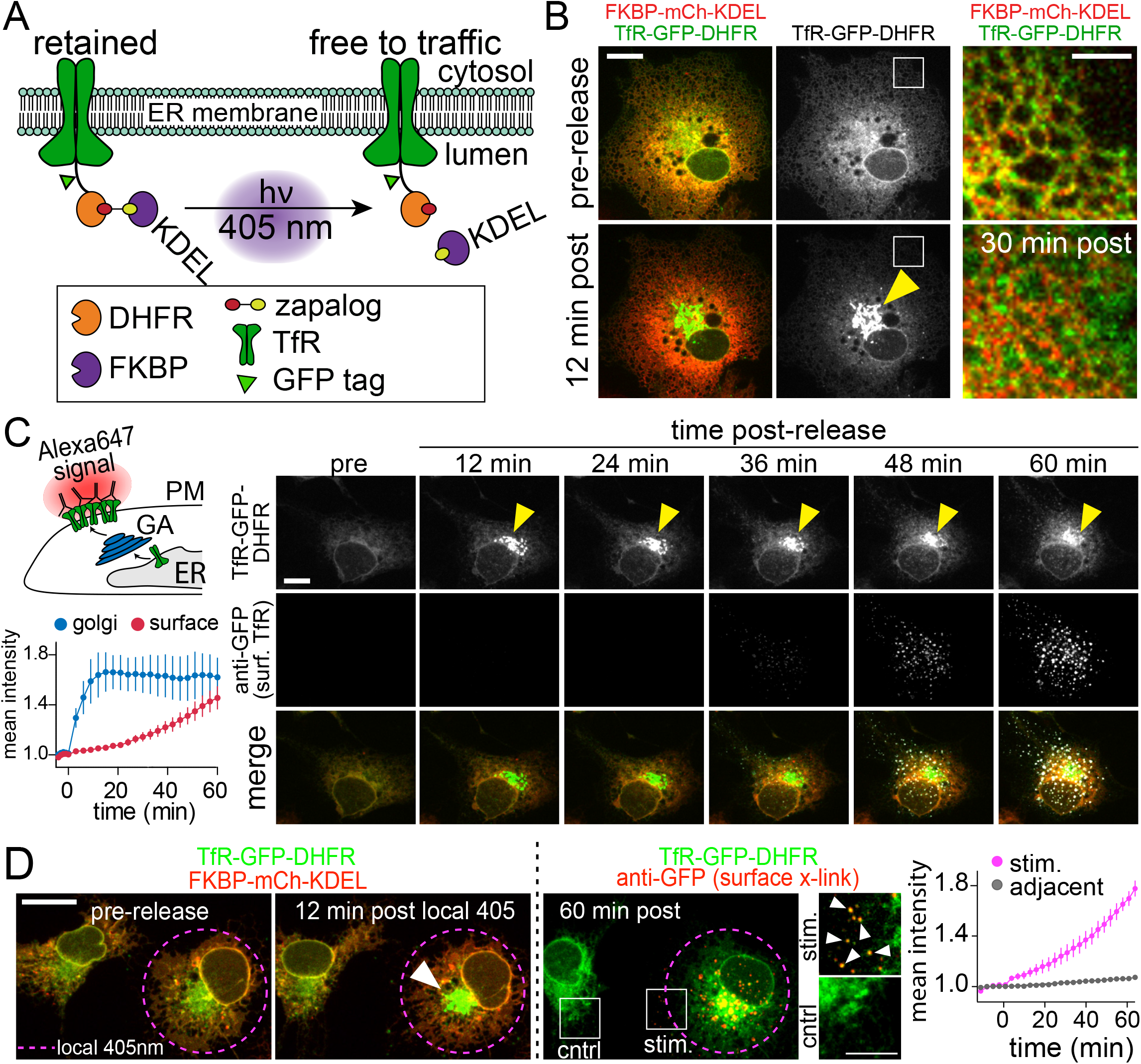
Developing and validating an approach for light-triggered protein secretion. **A**. Schematic of zapERtrap strategy for light-triggered ER release. DHFR is fused to the lumenal domain of an integral membrane protein cargo molecule (TfR shown here). An ER-retained version of FKBP (bearing a C-terminal KDEL sequence) is expressed in the lumen of the ER. The zapalog compound bridges these domains, retaining the DHFR-cargo molecule (TfR-GFP-DHFR shown here) in the ER. 405 nm light disrupts the zapalog tether between DHFR and FKBP allowing forward trafficking to proceed. **B**. 405 nm excitation triggers redistribution of TfR-GFP-DHFR (green) from the ER to the GA (arrowhead) while the FKBP-mCh-KDEL signal (red) remains in the ER. Magnified images (taken from the region marked by the white boxes) before (*top*) and 30 min following (*bottom*) ER release are shown on the right. The yellow arrowhead denotes the GA. Scale bar, 20 μm. Inset scale bar, 5 μm. **C**. Live-cell antibody surface labelling reveals the kinetics and spatiotemporal dynamics of cargo surface accumulation following ER release. *Top Left:* Schematic of strategy for real-time visualization of surface accumulation using an Alexa Fluor 647-conjugated anti-GFP (Alexa647-anti-GFP), which rapidly binds to and crosslinks TfR-GFP-DHFR as it appears on the cell surface. *Right*: Image time series of TfR-GFP-DHFR accumulation in the GA (top panels; arrowhead) and on the cell surface (middle panels) following light-triggered ER release at time 0. Scale bar, 15 μm. *Bottom Left:* Kinetics of TfR-GFP-DHFR accumulation in the GA (blue) and on the cell surface (red) following release. Data are represented as mean ± SEM (n=6 cells from 2 independent experiments). **D**. Local 405 nm excitation triggers ER release of TfR-GFP-DHFR in individual COS7 cells. *Left:* Focal illumination with 405 nm light (region marked by the pink dashed line) triggers TfR-GFP-DHFR (green) trafficking to the GA (arrowhead) the surface (right, 647-anti-GFP puncta shown in red) only in the photoactivated cell. Magnified images (taken from the regions marked by the white boxes) to the right show TfR-GFP-DHFR surface accumulation (arrowheads) in the photoactivated cell (“stim.”; *top*) and neighboring control cell (*bottom*). Scale bar, 20 μm. Inset scale bar, 10 μm. The plot to the right shows the average time course of TfR-GFP-DHFR surface accumulation for photoactivated cells (pink line) and neighboring control cells (gray line) that were not photoactivated. Data are represented as mean ± SEM (n=6 photoactivated cells, 8 control cells from 2 independent experiments).

Due to its robust trafficking properties, we initially developed and validated the system using the type II integral membrane protein transferrin receptor (TfR), which predominantly localizes to recycling endosomes (REs). Using COS7 cells, we show that in the absence of zapalog, TfR-GFP-DHFR strongly colocalizes with RE markers Rab11 and DHHC2 confirming that the DHFR tag does not disrupt subcellular targeting (Fig. S1A-C). However, in the presence of 500 nM zapalog (added immediately following transfection) and co-expressed mCh-FKBP-KDEL, TfR-GFP-DHFR was strongly retained in the ER (Fig. 1B). Following brief (10-50 msec) full field illumination with 405 nm light we observed redistribution of TfR-GFP-DHFR from the ER to the perinuclear GA with an average time-to-peak of 15.5 ± 1.43 min (Fig. 1B,C, Movie S1), similar to previously reported ER to GA trafficking kinetics using different inducible ER release systems (Boncompain et al., 2012; Chen et al., 2013; Horton and Ehlers, 2003; Presley et al., 1997). To measure subsequent plasma membrane insertion, we performed real-time surface labeling using an Alexa Fluor 647-conjugated antibody against the engineered GFP tag (Alexa647-anti-GFP) (Fig. 1C). As the GFP-tagged cargo is presented on the cell surface, rapid antibody binding results in signal amplification as well as restriction of lateral mobility through crosslinking, thereby allowing us to monitor integrated TfR-GFP-DHFR surface delivery in real time with high sensitivity (Fig. 1C, Movie S1). Following release of ER-retained TfR-GFP-DHFR, surface-bound Alexa647-anti-GFP could be detected within 20 min, similar to previous reports for the latency between ER release and surface appearance, confirming the effectiveness of our live-cell labeling strategy (Chen et al., 2013; Presley et al., 1997) (Fig. 1C). The absence of appreciable surface TfR-GFP-DHFR labeling prior to release confirmed efficient ER retention prior to light exposure (Fig. 1C).

One of the major potential applications of this approach is local control of secretory trafficking from user-defined cell populations or even subcellular domains. To test if this was possible with zapERtrap, we initiated ER release in only one of the COS7 cells within the imaging field with focally directed 405 nm excitation (Fig. 1D). We observed accumulation of TfR-GFP-DHFR in the GA in photoactivated cells but not in adjacent control cells (Fig. 1D, Movie S1). Accordingly, only photoactivated cells accumulated surface TfR-GFP-DHFR (Fig. 1D). Together, these data indicate that the zapERtrap inducible ER release system can be used to rapidly and locally trigger forward secretory trafficking with light.

### Spatial and temporal properties of synaptic protein trafficking in hippocampal neurons

We next asked whether we could use zapERtrap to investigate the trafficking itineraries for synaptic proteins in neurons. We engineered DHFR-fusions with the AMPA-type glutamate receptor GluA1 (DHFR-mNeon/GFP-GluA1) and the synaptic cell adhesion molecule neuroligin 1 (DHFR-GFP-NL1) (Fig. 2A). We first confirmed that the DHFR tag does not perturb their normal enrichment at dendritic spines, the predominant postsynaptic compartments of excitatory synapses (Fig. S2A). We next demonstrated these cargoes could be retained in the neuronal ER with zapalog and efficiently released with a brief light exposure (Fig. 2A,B; Movies S2, 3). Following a single 50 ms pulse of 405 nm light, TfR-GFP-DHFR and DHFR-GFP-NL1 exited the ER and redistributed to the GA within minutes (time to peak: 10.5 ± 0.48 min [TfR], 11.3 ± 0.85 min [NL1]) (Fig. 2A-C). Surprisingly, we observed much slower GA accumulation of DHFR-mNeon-GluA1, with a time to peak of 26.5 ± 1.53 min. (Figs. 2A-C, Movie S3).

**Figure 2.**
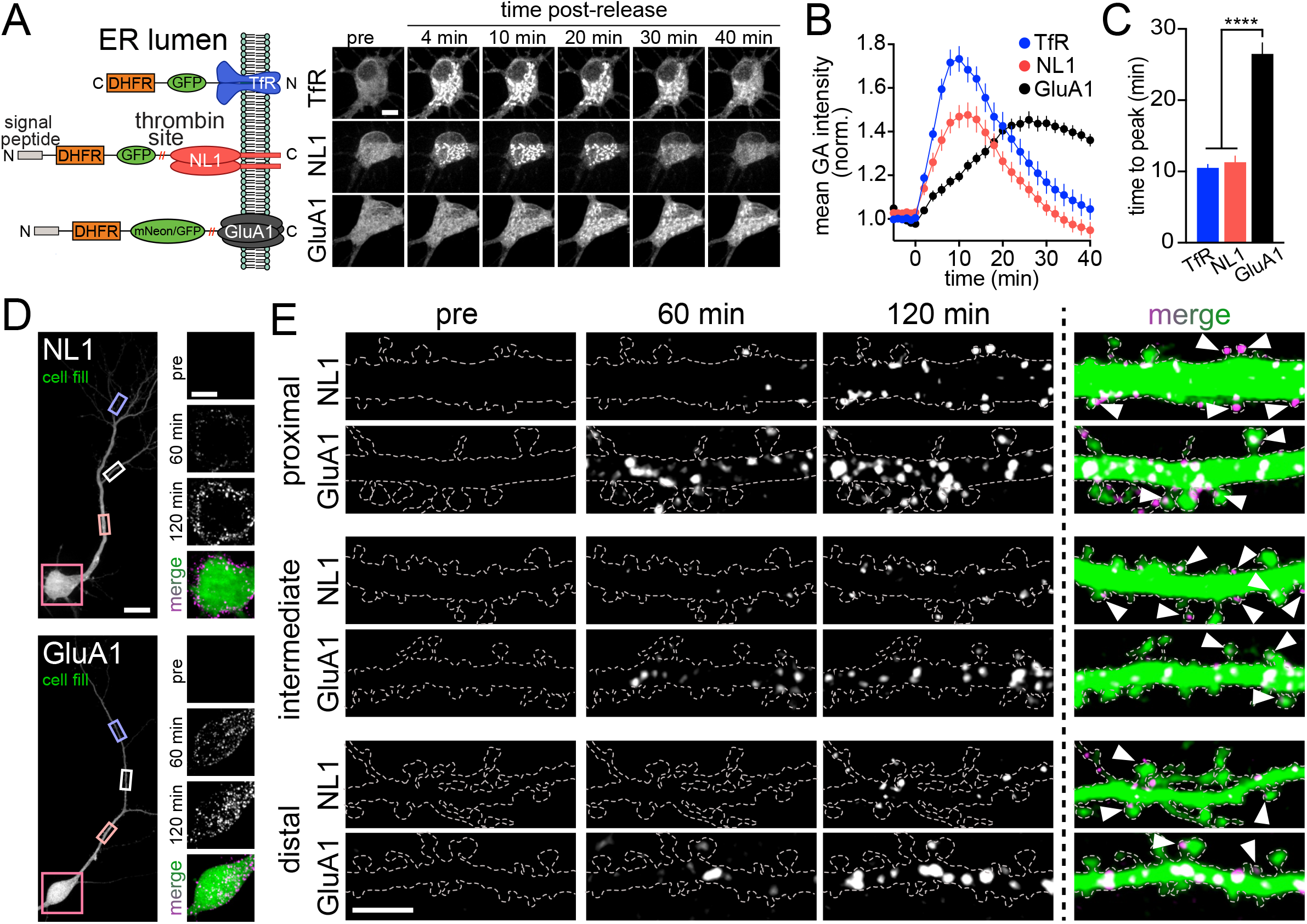
Spatial and temporal properties of synaptic protein trafficking in hippocampal neurons. **A**. *Left:* Schematic of constructs used for ER release experiments in hippocampal neurons. For simplicity, only one receptor subunit is depicted as the expression construct. *Right:* Image series of TfR-GFP-DHFR (*top*), DHFR-GFP-NL1 (*middle*) and DHFR-mNeon-GluA1 (*bottom*) accumulation in the GA following light-triggered ER release at time 0. Scale bar, 10 μm. **B**. Time course of TfR-GFP-DHFR (blue), DHFR-GFP-NL1 (red) and DHFR-mNeon-GluA1 (black) accumulation in the GA following ER release. Data are represented as mean ± SEM (n=11-27 cells from at least 2 independent experiments). **C**. Comparison of the time to peak accumulation for TfR-GFP-DHFR (blue), DHFR-GFP-NL1 (red) and DHFR-mNeon-GluA1 (black) following ER release. ****p<0.0001 (one-way ANOVA, Tukey’s multiple comparisons test). **D**. Localization of surface DHFR-GFP-NL1 (*top*) and DHFR-GFP-GluA1 (*bottom*) at the soma before, 60min and 120min following ER release (right panels). The insets are taken from the neuron at left (pink square). Scale bar, 20 μm. Inset scale bar, 10 μm. **E**. Localization of surface NL1 and GluA1 in the proximal (*top*), intermediate (*middle*) and distal (*bottom*) dendrites (taken from the peach, white and lavender rectangles in [B]) before, 60min and 120min following ER release. Arrowheads denote spines with surface label. Scale bar, 10 μm.

Given the large difference in the rate of NL1 and GluA1 accumulation at the GA, we asked whether there were also differences in the kinetics or location of cell surface accumulation. We first validated our surface labeling/crosslinking strategy in neurons (Figs. S2B-G). We quantified the rate of antibody binding by adding anti-GFP (directly conjugated to Alexa647) to live hippocampal neurons expressing DHFR-GFP-GluA1 that had not been ER retained. Antibody signal was detectable within seconds and accumulated with kinetics that were well approximated with a single exponential fit (*τ*=1.66 min) (Fig. S2B). We next confirmed that anti-GFP signal accumulating over longer time periods following ER release was indeed trapped on the cell surface, and did not arise from surface cargo that had bound antibody prior to being recaptured into intracellular compartments. We added an Alexa568-conjugated anti-rabbit secondary antibody to the extracellular solution of live cells that had previously accumulated DHFR-GFP-GluA1/Alexa647-anti-GFP (rabbit) for 80 min following ER release (Fig. S2F). Nearly all (86.6 ± 2.83 %) of the accumulated anti-GFP signal co-labeled with the anti-rabbit antibody signal, confirming surface localization (Fig. S2F). Finally, to verify surface label was effectively crosslinked and therefore immobilized in the general vicinity of surface delivery, we performed fluorescence recovery after photobleaching (FRAP) measurements (Fig. S2G). We observed nearly no fluorescence recovery when we photobleached accumulated Alexa647-anti-GFP signal over small dendritic segments, indicating effective immobilization of surface cargo (Fig. S2G).

With the labeling strategy validated, we next compared the spatial and temporal dynamics of NL1 and GluA1 surface trafficking following global (i.e. the entire cell was exposed to 405nm light) ER release (Fig. 2D-E). Hippocampal neurons expressing DHFR-GFP-GluA1 or DHFR-GFP-NL1 were imaged in extracellular solution containing Alexa647-anti-GFP during a 5-10 min baseline period, followed by exposure to a single, full field 405 nm light exposure. Cells were then imaged for an additional 120 min to monitor the kinetics and spatial distribution of surface accumulation (Figs. 2D,E; 3; Movie S4). For both NL1 and GluA1 we observed anti-GFP signal abruptly appear as discrete and stable punctate structures in the soma and dendrites, with the first detectable signal appearing approximately 20 min following ER release (Figs. 2D,E; S2C-E Movie S4). At 120 min post-release, both cargoes exhibited surface accumulation in the soma and proximal regions of the dendritic arbor (Figs. 2D,E). However, their dendritic distribution appeared markedly different. For GluA1, signal was easily discernable in more distal (>40 μm from the soma) dendrites compared to NL1, which appeared more enriched on the cell body compared to dendrites (Figs. 2D,E). To more precisely quantify the spatial profile of surface delivery, we computationally segmented discrete surface accumulations and classified whether they occurred in the soma, proximal (0-40μm from the soma) or distal (40-200μm from the soma) dendritic compartments (Figs. 3A-D; Movie S5). Surprisingly, even though GA accumulation was nearly 2.5-fold slower for GluA1 compared to NL1 (Fig. 2C), surface insertion within the somatic region was not significantly different (time to 10% of total surface accumulation: 50.0 ± 2.29 min [NL1], 41.5 ± 3.81 min [GluA1]) (Fig. 3A,B,D). For proximal dendritic regions (largely encompassing basal dendrites and the proximal region of the apical dendrite), the rates of surface accumulation were also similar between the two cargoes (time to 10% surface accumulation: 52.8 ± 3.07 min [NL1], 46.9 ± 4.03 min [GluA1]) (Figs. 2E; 3A,B). However, in more distal dendrites (dendritic regions that are >40 μm from the soma), GluA1 could be detected significantly sooner following ER release compared to NL1 (time to 10% surface accumulation: 66.6 ± 3.18 min [NL1], 50.9 ± 3.14 min [GluA1]) (Fig. 3A-D). Thus, GluA1 traffics to the surface more uniformly across the dendrites and soma while NL1 is delivered in a wave-like manner, with early insertion occurring mainly in the soma and proximal dendrites followed by more delayed insertion at distal dendrites (Fig. 3D; Movie S5).

**Figure 3.**
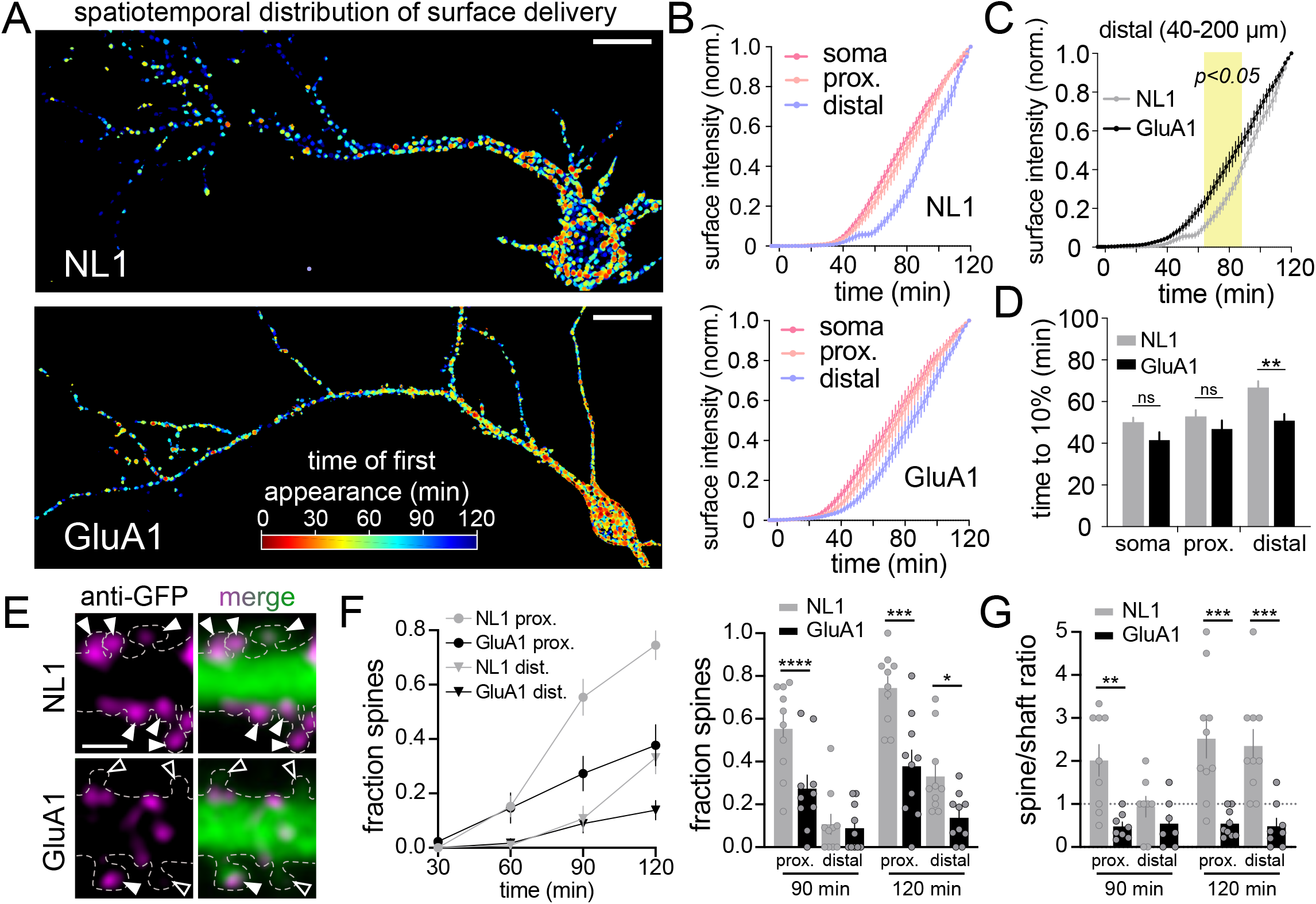
Subcellular distribution of NL1 and GluA1 surface presentation following global ER release. **A**. The timing and location of surface signal for the NL1-expressing (*top*) and GluA1-expressing (*bottom*) neurons are shown. Surface signal with a shorter latency of appearance following ER release are rendered in warmer colors. Scale bars, 20 μm. **B**. Time course of NL1 (*top*) and GluA1 (*bottom*) surface accumulation following ER release. Shown are the normalized intensities for surface signal at the soma (pink line), proximal regions of the dendritic arbor (peach line; up to 40 μm from the soma) or distal regions of the dendritic arbor (lavender line; 40-200 μm from the soma). Data are represented as mean ± SEM (NL1: n=10 neurons/timepoint from 4 independent experiments; GluA1: n=11 neurons/timepoint from 2 independent experiments). **C**. Comparison of NL1 (gray) and GluA1 (black) surface accumulation occurring at the distal regions of the dendritic arbor (40-200 μm from the soma) following ER release. Data are represented as mean ± SEM. The values within the yellow shaded region are significantly different from one another; p<0.05, two-way ANOVA, Bonferroni’s multiple comparisons test. **D**. Time to 10% of the total surface accumulation (measured at 120 min) is plotted for NL1 (gray) and GluA1 (black) for different cellular domains. Data are represented as mean ± SEM. **p<0.01, ns=not significant (unpaired t test). **E**. Representative images of NL1 (*top*) and GluA1 (*bottom*) surface signal in spines 90 min following ER release. Solid arrowheads denote cargo-positive spines. Open arrowheads mark spines that lack detectable surface cargo. The outline of the cell is shown as a dashed line and was drawn based on the GFP fill. Scale bar, 2 μm. **F**. Time course plotting the fraction of spines in proximal (circles) and distal (triangles) dendrites that contain surface NL1 (gray) or GluA1 (black) following ER release. A comparison of the fraction of NL1- and GluA1-positive spines at 90 min and 120 min is shown on the right. Data are represented as mean ± SEM. ****p<0.0001, ***p <0.001, *p<0.05 (unpaired t test; n=10 neurons/timepoint for NL1 and GluA1). **G**. Spine to shaft ratios of surface signal for NL1 (gray) or GluA1 (black) in proximal and distal dendrites 90 min and 120 min following ER release. Data are represented as mean ± SEM. ***p<0.001, **p<0.01 (unpaired t test; n=8-10 neurons/timepoint [NL1], n=7-9 neurons/timepoint [GluA1]).

We next asked whether there were any differences between GluA1 and NL1 accumulation at the surface of dendritic spines, the primary postsynaptic compartments of excitatory synapses. While the surface labeling method cannot distinguish between direct insertion into spines and lateral diffusion into spines prior to antibody crosslinking and detection, we observed large differences in the fraction of spines with accumulated signal. In comparison to GluA1, NL1 was targeted to a higher fraction of spines and more of the total dendritic signal resided in spines, compared to GluA1 (Figs. 3E-G). Thus new proteins can be rapidly incorporated into spines shortly following surface presentation, although the extent of spine localization appears to be highly cargo dependent.

### The axon initial segment is a surface trafficking hot spot for select cargoes

In addition to somatic and dendritic delivery, we also observed striking surface insertion of NL1 at the axon initial segment (AIS) very early following ER release (Fig. 4A-E; Movie S6). To ensure the signal we observed did not arise from intracellular vesicles loaded with labeled NL1 that had been at the surface elsewhere in the cell, we added an Alexa-568 anti-rabbit antibody (which binds the Alexa647-labeled anti-GFP used to detect cargo delivery) to live cells that had accumulated anti-GFP signal for 50 min following ER release. We observed nearly complete overlap between the anti-GFP signal and the anti-rabbit secondary, confirming AIS surface localization of newly trafficked NL1 (Fig. S3A). We also wanted to rule out the possibility that AIS insertion was due to overloading the secretory pathway, causing NL1 to “spill over” into a non-relevant, AIS-directed pathway. Here we leveraged the ability to precisely titrate the amount of NL1 released from the ER simply by decreasing the intensity of the excitation light, a major advantage of zapERtrap compared to other ER retention/release tools. We calibrated the amount of NL1 release by quantifying its accumulation in the GA following excitation with different light intensities (Fig. 4F). Importantly, we still observed strong surface delivery at the AIS, even at sub-saturating excitation intensities that released much less NL1, down to the threshold for reliably detecting subsequent surface accumulation (Figs. 4F,G). Surprisingly, we observed very little GluA1 signal at the AIS even at saturating light intensities that released the maximum level of GluA1 from the ER (Figs. 4B,G). Since the AIS is a major site of action potential initiation, we also tested whether NL1 trafficking at the AIS is regulated by neuron firing. We observed no difference in NL1 AIS accumulation in the presence of the voltage-gated sodium channel blocker tetrodotoxin (TTX) which suppresses action potentials or the GABA_A_ receptor antagonist bicuculline (Bic), which causes increased overall network activity and action potential firing (Fig. 4H). Combined, these data reveal a novel trafficking route to the surface of the AIS that is independent of neural activity, but highly cargo selective. We note that we were able to detect AIS enrichment in our experiments because our detection strategy effectively traps and immobilizes cargo soon after it reaches the cell surface, thus integrating the accumulated AIS signal. Indeed, if we delayed the addition of our immobilizing/detection antibody to 3h post ER release (when nearly all of the released NL1 had already trafficked to the surface) we did not observe any enrichment at the AIS. Thus, under normal steady-state conditions, NL1 that is initially presented at the AIS is efficiently recaptured by endocytosis or exits the AIS via lateral diffusion and trafficked elsewhere in the cell (Fig. S3B).

**Figure 4.**
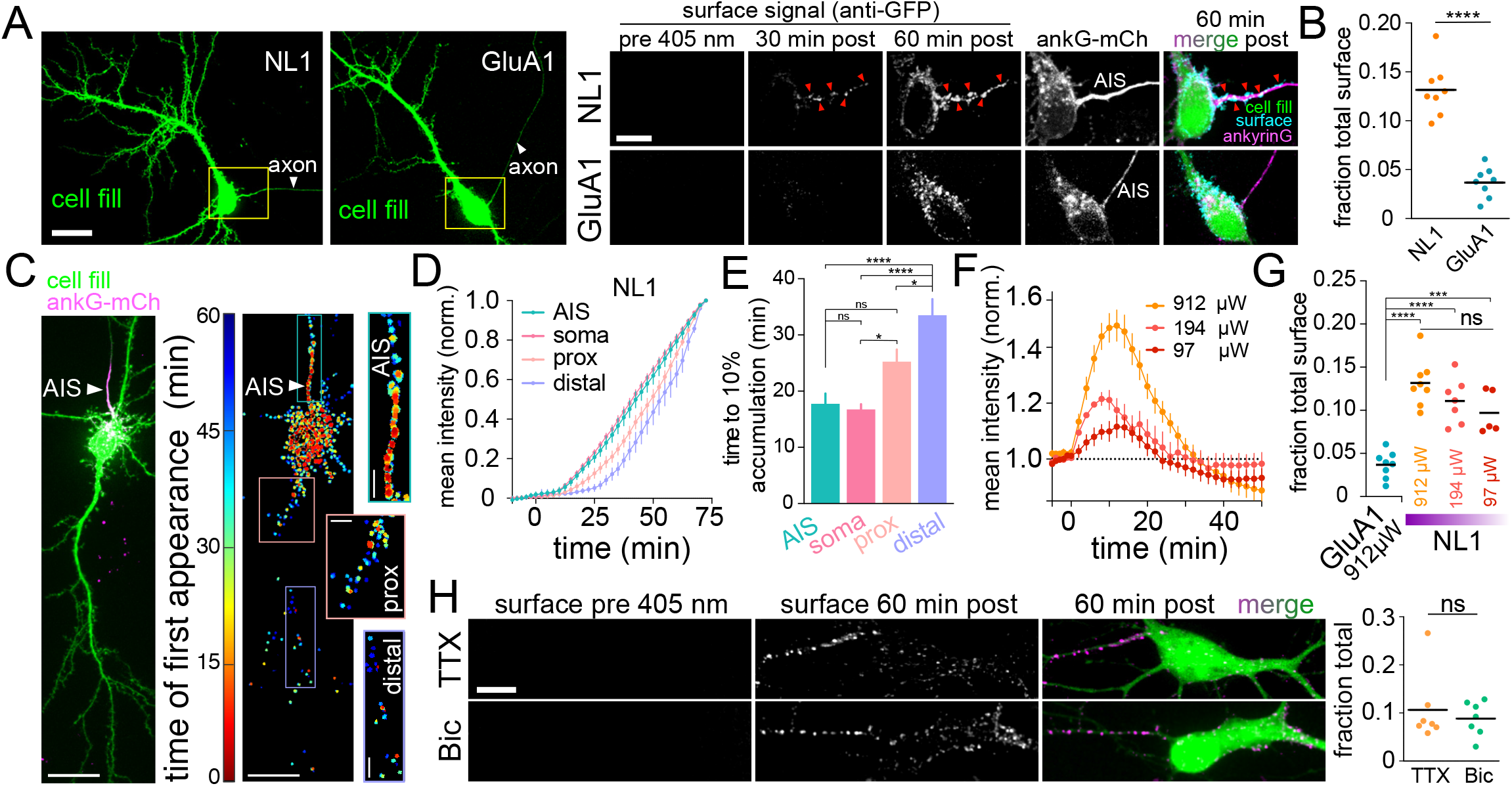
The axon initial segment is a surface trafficking hot spot for select cargoes. **A**. Representative images of NL1- and GluA1-expressing neurons co-transfected with the AIS marker ankyrinG-mCh (not shown) and a cell fill (green). The axon is indicated by the white arrowhead. Insets (taken from the yellow boxes) of NL1 (*top*) and GluA1 (*bottom*) surface accumulation at the AIS surface PM before and 60 min after ER release are shown on the right. The merged images show the surface accumulation (cyan) at the AIS marked by ankyrinG-mCh (magenta) 60 min post release. Red arrowheads denote surface cargo accumulations at the AIS PM. Scale bar, 20 μm. Inset scale bar, 10 μm. **B**. Quantification of AIS surface signal (expressed as a fraction of total surface signal) for NL1 (orange) or GluA1 (blue) 60 min following ER release. Data are represented as mean ± SEM. ****p<0.0001 (unpaired t test, n=8 from 2 independent experiments for NL1 and GluA1). **C**. Confocal image (*left*) and map showing the timing and location of NL1 surface appearance at the AIS (*right*). The AIS marker ankyrinG-mCh (magenta) was used to confirm AIS identity. Scale bars, 20 μm. Insets showing the AIS, proximal and distal dendrites (taken from the teal, peach and lavender boxes) highlight the spatiotemporal differences in surface NL1 accumulation. Scale bars, 20 μm. Inset scale bars, 5 μm. **D**. Time course of plotting NL1 surface signal at four distinct subcellular locations. Shown are the mean NL1 surface intensities (normalized to their maximum values) at the AIS (teal line), soma (pink line), proximal dendrites (5-40 μm from the soma; peach line) or distal dendrites (40-200 μm from the soma; lavender line). Data are represented as mean ± SEM (n=10 neurons/timepoint from 2 independent experiments). **E**. Time taken to for surface NL1 signal to reach 10% of maximum in each subcellular compartment. Data are represented as mean ± SEM. *p<0.05, ****p<0.0001, ns=not significant (one-way ANOVA, Tukey’s multiple comparisons test; n=10 neurons from 2 independent experiments). **F**. NL1 release from the ER can be titrated. Shown is the quantification of NL1 GA accumulation following illumination with different 405nm light intensities (912 μW, 194 μW and 97 μW). Data are represented as mean ± SEM. (n=7-14 neurons/condition from at least 2 independent experiments). **G**. NL1 still accumulates in the AIS when minimal NL1 is released from the ER. Quantification of NL1 signal at the AIS (expressed as a fraction of the total signal for each cell), 60 min post ER release following exposure to different light intensities. GluA1 surface signal at the AIS using a saturating light intensity (912μW) is shown for comparison. Data are represented as mean ± SEM. ***p<0.001, ****p<0.0001, ns=not significant (one-way ANOVA, Tukey’s multiple comparisons test; n=5-8 neurons/condition from 2 independent experiments). **H**. Neural activity (suppressed with TTX, top; elevated with Bic, bottom) does not influence NL1 trafficking to the surface of the AIS. Scale bar, 10 μm. Quantification of NL1 signal at the AIS surface 60 min following ER release in the presence of TTX (orange) or Bic (green) is shown on the right. Data are represented as mean ± SEM. ns=not significant (unpaired t test; n=7 neurons/condition from 3 independent experiments).

### Rate, spatial dynamics and activity-dependence of secretory trafficking from dendritic compartments

We next assessed the degree to which the subcellular site of ER exit dictates where proteins end up on the neuronal cell surface. A major feature of zapERtrap is that it allows local ER release from user-defined subcellular regions. We first tested whether protein cargoes exiting the ER within a specific dendritic branch are selectively delivered to the nearby plasma membrane within the same branch, or are dispersed as they move through mobile downstream trafficking organelles. We also asked whether neural activity influences the spatial pattern of surface expression by either suppressing or stimulating neuronal firing by applying TTX or Bic prior to ER release (Fig. 5A). We locally released GluA1 and NL1 from dendrites over a region approximately 30-40 μm in length and 30-50μm from the soma). Following release, we observed punctate structures form within tens of seconds in the illuminated dendrite but not in neighboring dendrites or the soma, consistent with a previous study demonstrating spatially restricted ER exit and ERGIC accumulation in dendrites (Figs. 5B, S4A-C) (Bowen et al., 2017). These structures were initially stable before abruptly moving away from their site of appearance, often rapidly leaving the photo-targeted region (Fig. S4D, Movie S7). Accordingly, we observed widespread surface delivery of released cargoes not only at the targeted dendrite but also at untargeted control dendrites and even the cell body, especially for NL1 (Figs. 5C,D). Under no condition could we detect a specific enrichment of overall surface signal within or near the illuminated region (Figs. 5E,F,G). Despite no difference in the overall amount of GluA1 delivered to the surface of the targeted dendrite relative to untargeted neighboring control dendrites, we found that spines within the illuminated region were significantly enriched with GluA1 compared to similarly sized regions in neighboring control dendrites, but only when action potentials were suppressed with TTX (Fig. 5H). In contrast, NL1 was delivered to a similar fraction of spines both inside and outside of the ER release zone, with no significant dependence on activity (Fig. 5H).

**Figure 5.**
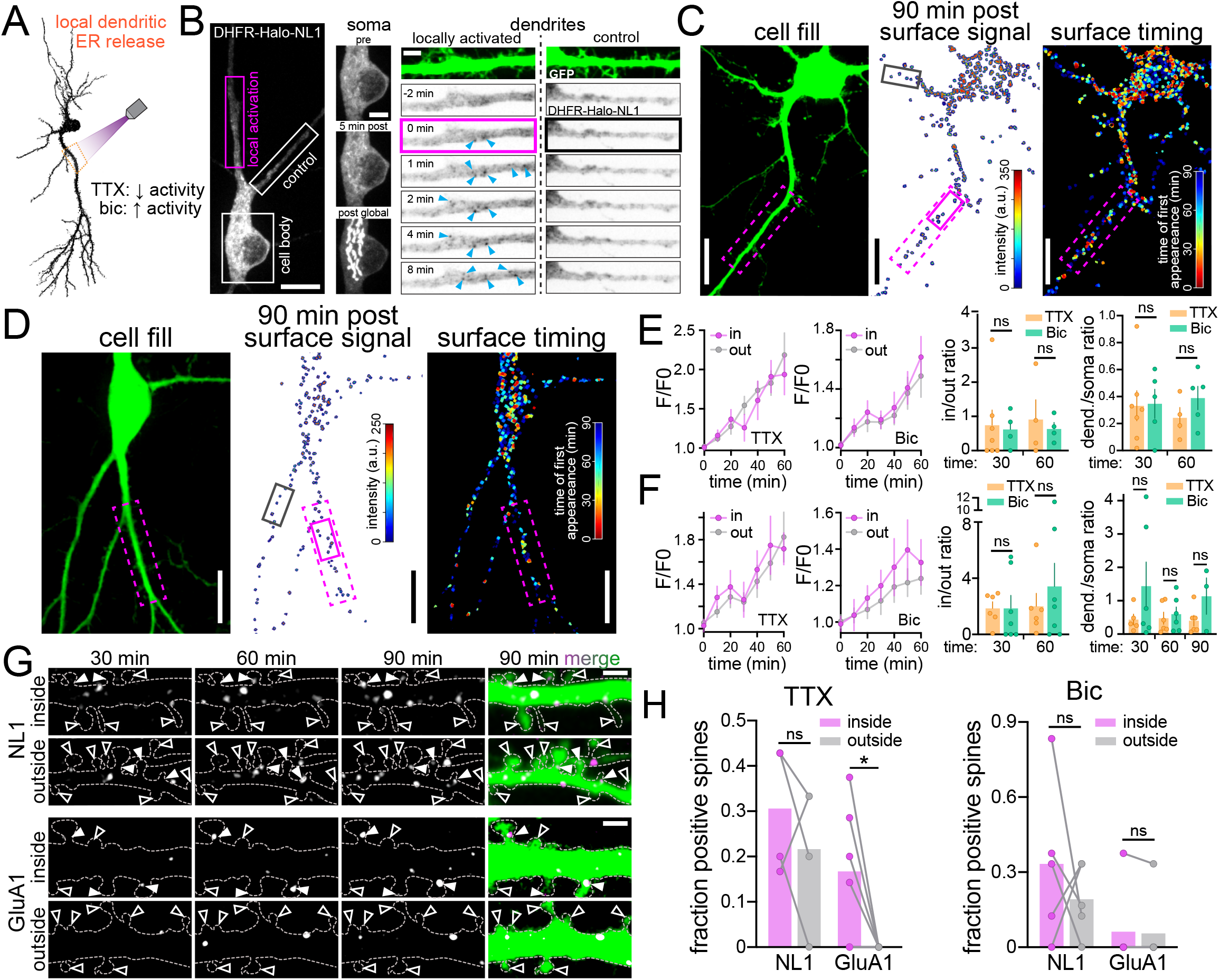
Rate, spatial dynamics and activity-dependence of secretory trafficking from dendritic compartments. **A**. Schematic of experimental strategy. NL1 and GluA1 were locally released from a defined region of a dendritic branch in the presence of either TTX to suppress neuronal activity or Bic to elevate neuronal activity. **B**. Representative image (*left*) and image time series of DHFR-HaloTag-NL1 subcellular distribution at the soma (*center*) and dendrites (*right*) shortly following local dendritic ER release (note that halotag was used in these experiments for superior detection of early intracellular trafficking events). Scale bar, 10 μm. The area indicated by the pink box was subject to 405 nm illumination and is shown to the right. Blue arrowheads mark appearance of vesicular structures shortly following illumination. The neighboring unstimulated control branch (white box in left image) is shown for comparison. At the end of the experiment the cell was exposed to global full field illumination and imaged 10 min later. Note the strong NL1 accumulation in the somatic GA following global but not local dendritic release (middle panels). All inset scale bars, 5 μm. **C**. Confocal image showing the cell fill (*left*), surface signal (*center*) and spatiotemporal map (*right*) for NL1 following local dendritic ER release. The photoactivated region is denoted by the dashed pink line. The black and pink (solid lines) boxes denote the dendritic regions shown in (G). Scale bars, 10 μm. **D**. Confocal image showing the cell fill (*left*), surface signal (*center*) and spatiotemporal map (*right*) for GluA1 following local dendritic ER release. The photoactivated region is denoted by the dashed pink line. The black and pink (solid lines) boxes denote the dendritic regions shown in (G). Scale bars, 10 μm. **E**. Timecourse of NL1 surface signal appearing within the dendritic ER release zone (purple) vs. dendrites up to 200 μm from the soma outside the release zone (gray) in the presence of TTX (*left*) or Bic (*right*). No significant differences were found at any timepoint (two-way ANOVA, Bonferroni’s multiple comparisons test). Data are represented as mean ± SEM. To the right is plotted the ratio of signal within the release zone vs outside the release zone (in/out) and the ratio of total dendritic to somatic signal (dend./soma ratio) 30 and 60 min following dendritic ER release in the presence of TTX (orange) or Bic (green). Data are represented as mean ± SEM. ns=not significant (unpaired t test; n=4-7 from at least 3 independent experiments. **F**. Timecourse of GluA1 surface signal appearing within the dendritic ER release zone (purple) vs. dendrites up to 200 μm from the soma outside the release zone (gray) in the presence of TTX (*left*) or Bic (*right*). No significant differences were found at any timepoint (two-way ANOVA, Bonferroni’s multiple comparisons test). Data are represented as mean ± SEM. To the right is plotted the ratio of signal within the release zone vs outside the release zone (in/out) and the ratio of total dendritic to somatic signal (dend./soma ratio) 30 and 60 min following dendritic ER release in the presence of TTX (orange) or Bic (green). Data are represented as mean ± SEM. ns=not significant (unpaired t test; n=3-7 from at least 2 independent experiments. **G**. Image series of NL1 (top panels) and GluA1 (bottom panels) surface accumulation inside vs. outside the photoactivated region depicted by the white and pink (solid) boxes in (C and D). Solid arrowheads denote cargo-positive spines. Open arrowheads mark spines that lack detectable surface cargo. The dashed lines show cell morphology and were drawn based on a cell fill (green signal in merge). Scale bars, 2 μm. **H**. Comparison of the fraction of cargo-positive spines within the release zone (purple) vs. randomly selected regions of the same size in separate control dendrites (gray) 60 min following local dendritic ER release in the presence of TTX (*left*) or Bic (*right*). Data are represented as mean ± SEM. *p<0.05, ns=not significant (paired t test; n=4-6 from at least 3 independent experiments.

### Rate, spatial dynamics and activity-dependence of secretory cargoes derived from the cell body

The extent to which cell body-derived proteins can be directly trafficked to the surface of dendrites through long-range vesicular transport remains unsettled (Adesnik et al., 2005; Hanus et al., 2014; Horton and Ehlers, 2003; Williams et al., 2016). To address this issue, we used focal illumination to specifically release target proteins from the somatic ER and asked whether they appeared on the surface of the soma, dendrites (or both) and whether neural activity influences the spatiotemporal dynamics of forward trafficking from the somatic compartment. We released NL1 and GluA1 specifically from the cell body ER in the presence of either TTX or Bic (Fig. 6A). Following local somatic release, cargo rapidly accumulated in the cell body GA with no detectable accumulation occurring in ERGIC or GA-like structures in either proximal or distal dendrites (Fig. 6B). Thus, cargoes released from the soma utilize the cell body secretory network rather than diffusing to dendritic ER exit sites (Cui-Wang et al, 2012; Aridor and Fish, 2009). In the minutes following somatic ER release, both NL1 and GluA1 appeared at the cell surface at the soma as well as tens to hundreds of micrometers into the dendritic arbor (Figs. 6C; S5A,B; Movie S8). NL1 also rapidly appeared at the surface of the AIS, similar to our global release experiments (Fig. 6C). Intriguingly, elevating neural activity with Bic increased the total amount of GluA1 delivered to the cell surface but did not have a significant effect on the timing of surface delivery at the soma or dendrites (Fig. 6D,F,G). Conversely, elevating activity did not impact the total amount of NL1 surface delivery but significantly decreased the time to achieve detectable surface signal at distal dendrites (Fig. 6D,E). Together these data support the presence of distinct trafficking networks for the two cargoes, even though both originated from the soma.

**Figure 6.**
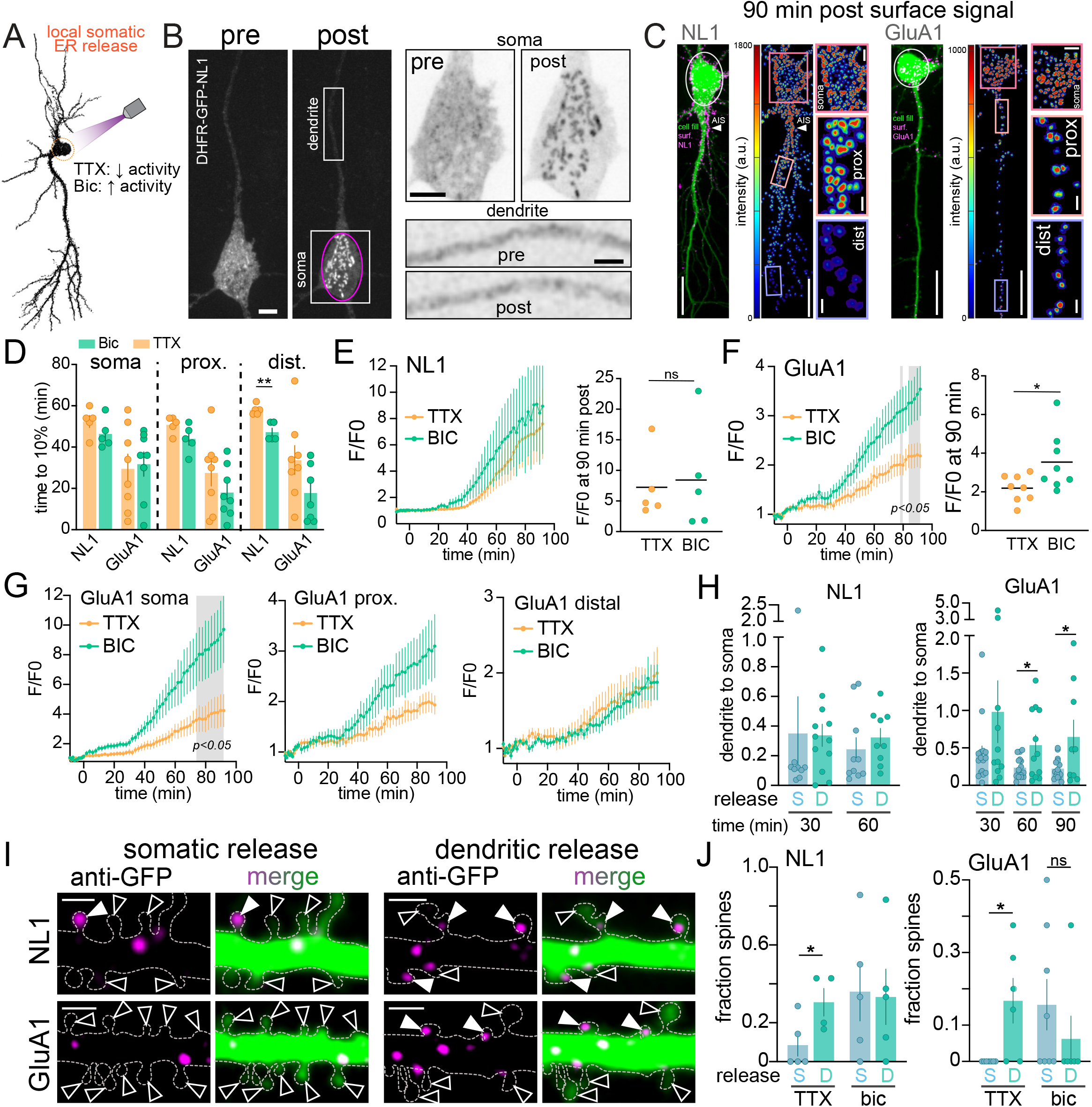
Rate, spatial dynamics and activity-dependence of secretory trafficking derived from the cell body. **A**. Schematic of experimental strategy. NL1 and GluA1 were locally released from the soma in the presence of either TTX to suppress neuronal activity or Bic to elevate neuronal activity. **B**. Representative example of DHFR-GFP-NL1 subcellular localization before (*left*) and 14 min after (*right*) somatic ER release. The area indicated by the pink circle was targeted with 405 nm illumination. The white boxes correspond to the inset images (shown on the right) of the soma (*top panels*) and dendrite (*bottom panels*) pre and post-ER release. Note the absence of any vesicular structures appearing in the dendrites following local somatic release. Scale bar, 6 μm. Top inset scale bar, 6 μm. Bottom inset scale bar, 3 μm. **C**. Merged confocal images showing cell fill (green) and surface signal (magenta) for NL1 (left) and GluA1 (right) 90min following ER release. Surface signal is shown to the right with warmer colors indicating higher intensity. The photoactivated regions are denoted by white circles. Insets of the soma, proximal dendrites and distal dendrites (taken from the pink, peach and lavender boxes) 90 min post-release. Scale bars, 20 μm. Soma inset scale bars, 5 μm. Dendrite inset scale bars, 2 μm. **D**. Time to 10% surface accumulation following somatic ER release in the presence of TTX (orange bars) or Bic (green bars) for NL1 and GluA1. Data are represented as mean ± SEM. **p<0.01, *p<0.05 (unpaired t test; n=5-8 neurons from at least 3 independent experiments) **E**. Time course of NL1 whole-cell fluorescence intensity following somatic ER release in the presence of TTX (orange line) or Bic (green line). Whole-cell fluorescence intensity at 90 min is shown on the right. Data are represented as mean ± SEM. ns=not significant (unpaired t test; n=5 neurons/condition from 3 independent experiments). **F**. Time course of GluA1 whole-cell fluorescence intensity following somatic ER release in the presence of TTX (orange line) or Bic (green line). The intensity values at the timepoints within the gray shaded regions are significantly different (p<0.05, two-way ANOVA, Bonferroni’s multiple comparisons test; n=8 neurons/condition from 2 independent experiments). A comparison of whole-cell fluorescence intensity at 90 min is shown on the right. Data are represented as mean ± SEM. *p<0.05 (unpaired t test). **G**. Time course of GluA1 surface trafficking at the soma (*left*), proximal dendrites (up to 40 μm from the soma; *center*) and distal dendrites (40-200 μm from the soma; *right*). Data are represented as mean ± SEM. The intensity values at the timepoints within the gray shaded region are significantly different (at least p<0.05, two-way ANOVA, Bonferroni’s multiple comparisons test). **H**. The ratio of total dendritic signal to total somatic signal is plotted at different timepoints following local somatic ER release (blue) or local dendritic ER release (teal) for NL1 (*left*) and GluA1 (*right*). Data are represented as mean ± SEM. *p<0.05 (unpaired t test; n=9-16 neurons from at least 2 independent experiments). **I**. Representative images of NL1 (*top*) and GluA1 (*bottom*) surface accumulation in the distal dendrites 60 min following local somatic (*left*) or local dendritic (*right*) ER release. The merged images show the cell fill (green) and Alexa647-anti-GFP signal (magenta). Solid arrowheads denote surface-cargo containing spines. Open arrowheads mark spines lacking detectable surface cargo. Scale bars, 2 μm. **J**. Comparison of the fraction of cargo-positive spines in the distal dendrites at 60 min following local somatic release (blue) vs. local dendritic release (teal) of NL1 (*left*) or GluA1 (*right*) in the presence of TTX or Bic. The values shown for dendritic ER release are measurements made from inside the targeted release zones. Data are represented as mean ± SEM. *p<0.05, ns=not significant (unpaired t test; n=4-8 neurons from at least 2 independent experiments).

**Figure 7.**
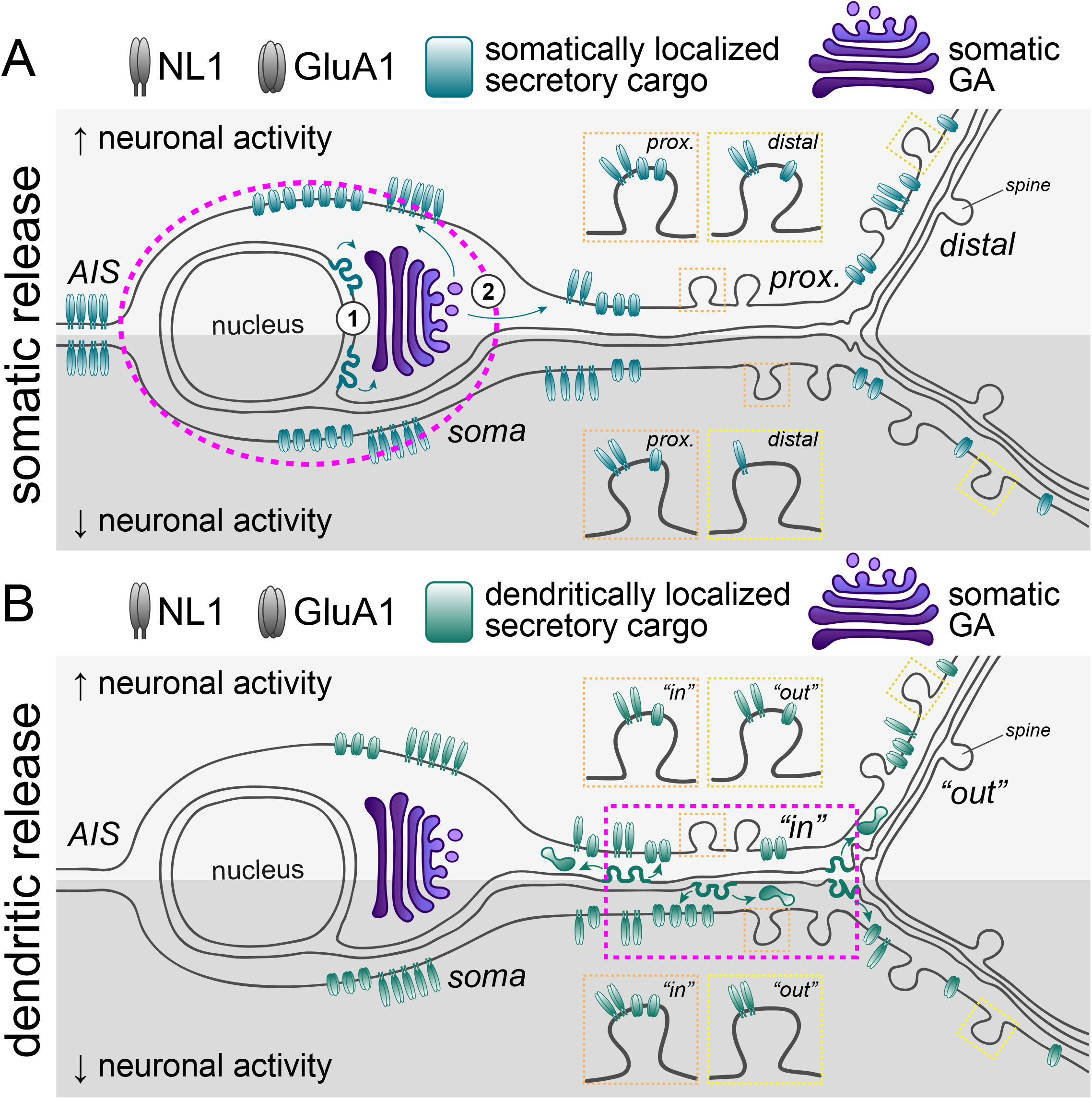
Models for subcellular secretory trafficking routes. Schematics highlighting cargo-specific, activity-dependent differences in the spatiotemporal dynamics of forward trafficking from the somatic (A) and dendritic (B) compartments. The pink dashed line marks the region of ER exit. **A**. Cargoes entering the somatic secretory network (1) traffic to the somatic GA and (2) appear at the surface at the soma and tens to hundreds of micrometers into the dendritic arbor. Elevated activity (lighter gray shaded region) impacts the timing of NL1 surface delivery (appears sooner at distal dendrites) without affect the total amount, whereas elevated activity impacts the total amount of GluA1 delivered to the surface. Both NL1 and GluA1 become more spine enriched with elevated activity. **B**. Cargoes entering the dendritic secretory network are rapidly dispersed before surface insertion in both high and low activity conditions. While overall surface presentation is not enriched near the ER release zone, local synapses are preferentially targeted when activity is suppressed.

Intriguingly, we also observed surface accumulation of both GluA1 and NL1 in dendritic spines, indicating that somatically derived proteins undergo long-range trafficking at or near postsynaptic compartments within tens of minutes. Spine targeting of both GluA1 and NL1 occurred in a proximal (higher fraction) to distal (lower fraction) gradient and was highly sensitive to neural activity, with elevated activity significantly increasing the overall fraction of cargo-positive spines (Fig. S5C-F).

While NL1 and GluA1 originating from either the somatic or dendritic secretory networks could be delivered to dendrites and spines, we noted some important differences between somatically and dendritically processed proteins. First, the dendrite/soma ratio of surface GluA1 (but not NL1) was significantly higher when GluA1 was released from dendrites, supporting a more highly compartmentalized local trafficking network for GluA1 (Fig. 6H). Second, accumulation within dendritic spines was much higher for dendritically vs. somatically released cargos, especially when activity was suppressed (Fig. 6I,J). Finally, elevating neural activity promoted somatic GluA1 trafficking to dendritic spines but appeared to suppress local spine accumulation of GluA1 released from dendrites. Conversely, TTX nearly completely suppressed spine targeting of somatically released GluA1 but strongly promoted spine localization of dendritically processed GluA1 (Fig. 6I,J). Thus the location of ER exit strongly influences subcellular targeting with neural activity exerting opposite effects on the same protein originating from different cellular domains.

## Discussion

The degree to which the neuronal secretory network is organized to support long-range or local trafficking has remained largely unexplored due to the lack of suitable approaches for selectively initiating protein trafficking from distinct subcellular domains. This remains a central issue in neuronal cell biology because in addition to constitutive protein turnover and homeostasis, prominent models for synaptic function and diverse forms of plasticity rely on synthesis, trafficking and targeting of key receptor proteins to remote cellular domains (Ju et al., 2004; Mameli et al., 2007; Schanzenbächer et al., 2018; Schilström et al., 2006; Smith et al., 2005; Sutton and Schuman, 2006). Here we engineered a new approach allowing remote control of forward secretory trafficking from user-defined subcellular regions. Our experiments reveal several new trafficking principles. First, long-range transport from the soma can support protein demand at synaptic sites hundreds of micrometers into the dendritic arbor within biologically relevant timescales. Second, the dendritic secretory pathway can support trafficking to local synaptic sites under specific conditions, but generally results in unexpectedly widespread delivery to the cell surface. Third, the AIS plays a surprising role as a major trafficking hotspot for initial surface insertion of select proteins. Finally, neural activity plays a major role in sculpting the spatial distribution, synaptic targeting and overall level of surface expression, with distinct trafficking rules for the same protein cargo originating from different cellular domains.

### Distinct, cargo-specific trafficking networks originate from the same subcellular compartment

Previous studies support distinct biosynthetic trafficking routes for diverse postsynaptic proteins (Bowen et al., 2017; Gu et al., 2016; Hanus et al., 2016; Jeyifous et al., 2009). For example, a significant fraction of AMPA receptors (but not NL1) appear to use a non-conventional trafficking route that bypasses the GA (Bowen et al., 2017; Hanus et al., 2016). However, it remains unclear if utilization of different trafficking networks is dictated by the identity of the cargo molecule and/or the subcellular domain in which it is synthesized and enters the secretory network. Surprisingly, we observed several major differences between NL1 and GluA1 trafficking, even when both proteins were released from the somatic ER. First, GluA1 trafficking to the somatic GA was drastically slower and in some cells difficult to detect, even though it appeared at the cell surface with similar kinetics as NL1, consistent with an alternative trafficking route for at least a fraction of AMPA receptors (Bowen et al., 2017; Hanus et al., 2016). Second, increasing neural activity elevated GluA1 surface delivery but had no impact on the total amount of NL1 delivered to the cell surface, consistent with previous studies (Evans et al., 2017; Hangen et al., 2018). This is also consistent with a previous study showing that a significant fraction of AMPA receptors traffic through recycling endosomes, whose fusion with the dendritic PM is stimulated by activity (Bowen et al., 2017; Kennedy et al., 2010; Park et al., 2004). Third, a significant fraction of NL1 but not GluA1, was trafficked to the PM of the AIS. Previous studies demonstrate intracellular trafficking vesicles harboring somatodendritic proteins are prevented from axonal entry immediately proximal to, or within the AIS (Al-Bassam et al., 2012; Farías et al., 2015; Song et al., 2009). However, transport vesicle fusion with the AIS PM has escaped previous detection because under normal steady-state conditions, cargo proteins utilizing this route are not enriched at the AIS. While further experiments will be necessary to establish the functional significance, fusion of these carriers at the AIS could be a novel sorting step for separating co-mingled dendritic and axonal proteins for subsequent recapture and transport to their respective cellular domains. Finally, NL1 signal appeared to consistently accumulate on a larger fraction of dendritic spines compared to GluA1. Given our antibody crosslinking approach can only approximate the location of surface insertion, we cannot distinguish whether cargo was directly inserted into the dendritic spine membrane, or inserted into the nearby dendritic shaft followed by diffusion into spines prior to antibody crosslinking and detection. Unfortunately, faster detection probes (e.g. superecliptic pHluorin) were not sensitive enough to pinpoint the location of insertion of individual trafficking vesicles. Nevertheless, these experiments clearly demonstrate that cargoes originating in the somatic compartment can rapidly cover long distances in the dendritic arbor where they are delivered near synaptic sites. The molecular determinants that direct these and other cargoes to distinct secretory networks and the collection of organelles responsible for differential trafficking properties will be addressed in future studies.

### The dendritic secretory network generally supports diffuse surface expression

Unlike smaller conventional cells, whose secretory network can rapidly satisfy protein demand throughout the cell, neurons have broadly distributed organelle networks that are hypothesized to support both centralized and satellite trafficking at remote sites (Bowen et al., 2017; Chirillo et al., 2019; Cooney et al., 2002; Hanus et al., 2016; Horton and Ehlers, 2003; Mikhaylova et al., 2016; Spacek and Harris, 1997; Torre and Steward, 1996). For example, substantial evidence supports local synthesis of integral membrane proteins and secreted factors at remote sites in dendrites, including the presence of their mRNAs, ribosome-associated ER, ER exit sites and ERGICs (Cajigas et al., 2012; Cui-Wang et al., 2012; Wu et al., 2017). It should be noted that local synthesis is not a requirement for dendritic ER localization as somatically-derived secretory proteins can also populate dendritic ER through lateral diffusion or active transport prior to ER exit (Cui-Wang et al., 2012; Valenzuela et al., 2014). In either case, the untested assumption is that proteins exiting the dendritic ER are delivered to nearby synaptic sites through a local, autonomous secretory network. Often overlooked is how spatially precise surface targeting could be (Bowen et al., 2017; Horton and Ehlers, 2003; Mikhaylova et al., 2016). Indeed, our data show that proteins released from targeted regions of dendritic ER generally reach the cell surface over a much broader area than the targeted zone of ER exit, in most cases appearing on the surface of neighboring dendrites, the soma and even the AIS (for NL1). Surprisingly, the overall dendrite-to-soma ratio of accumulated NL1 surface signal was nearly indistinguishable when it was released from dendrites or the soma. This could be due to a significant fraction of dendritically released NL1 converging in a common transport organelle shared with the somatic network. It is also possible that a fraction of dendritically released NL1 is retrogradely directed to the somatic GA for central processing. Although we did not observe appreciable accumulation of intracellular NL1 at the somatic GA following dendritic release, it is possible that a small increase in GA signal was masked by the persistent signal from unreleased, ER-localized NL1 in the cell body. In contrast, the overall dendrite-to-soma ratio of GluA1 was higher following dendritic release, supporting a distinct and more highly compartmentalized trafficking network. Although overall surface trafficking for both cargoes was more widespread than expected, dendritically released NL1 and GluA1 were both targeted to a higher percentage of dendritic spines compared to their somatically processed counterparts. This could be due to differential posttranslational modifications of dendritic proteins that favor binding to synaptic sites, or local dendritic trafficking routes that directly supply specialized trafficking organelles in spines. Whether resident spine organelles, including the ER/spine apparatus, recycling endosomes and/or lysosomes, could act as direct conduits to specific spines for dendritically released cargoes await further investigation (Bowen et al., 2017; Chirillo et al., 2019; Goo et al., 2017; Kennedy et al., 2010; Kulik et al., 2019; Padamsey et al., 2016; Park et al., 2004; 2006; Spacek and Harris, 1997).

### Implications for local protein control during synaptic activity and plasticity

Whether the physical arrangement and functional properties of dendritic trafficking organelles could support the timing and spatial precision of local dendritic protein trafficking proposed in current models of synaptic function and plasticity has remained largely unexplored (Ju et al., 2004; Kang and Schuman, 1996; Linden, 1996; Mameli et al., 2007; Sutton and Schuman, 2006). While the activity manipulations in our experiments were relatively basic, we observed no clear evidence for precise local surface expression for proteins released from the dendritic ER, with one notable exception. Despite relatively diffuse overall surface expression, we found that GluA1 AMPA receptors specifically targeted dendritic spines near the targeted ER release zone when neural activity was suppressed with TTX. Intriguingly, the time course of surface expression and spine localization of dendritically processed GluA1 is consistent with previous studies demonstrating local GluA1 synthesis, surface expression and spine targeting during homeostatic synaptic strengthening initiated by prolonged suppression of neural activity and synaptic glutamate receptors (Aoto et al., 2008; Sutton and Schuman, 2006). Surprisingly, the opposite was true for AMPA receptors processed in the soma, highlighting a major difference in trafficking for the same protein originating from different subcellular domains.

More rapid and synapse-specific forms of plasticity such as long term potentiation (LTP) initially rely on existing pools of synaptic proteins that are driven to synaptic sites by lateral diffusion and/or surface trafficking through regulated fusion of recycling organelles (Hiester et al., 2017; Kennedy et al., 2010; Lledo et al., 1998; Makino and Malinow, 2009; Park et al., 2004; Patterson et al., 2010; Penn et al., 2017). Indeed, the kinetics of surface expression we observe for proteins released from the somatic or dendritic ER are much too slow to account for fast (within seconds to minutes) synaptic alterations mediated by newly synthesized membrane proteins. On the other hand, previous studies demonstrated that AMPA receptors are synthesized and that secretory trafficking is required for sustained synaptic potentiation following LTP induction (Broutman and Baudry, 2001; Ju et al., 2004; Kramár et al., 2012; Nayak et al., 1998). For example, pharmacologically disrupting secretory trafficking shortly following LTP induction led to a decline in potentiated synaptic responses to pre-induction baseline levels in ∼1 hour, similar to the time course of dendritic protein delivery from either somatic or dendritic secretory networks (Kramár et al., 2012). Future experiments combining diverse plasticity mechanisms with zapERtrap will be a powerful approach for addressing whether secretory cargoes can be more precisely targeted to individual dendritic branches or individual synapses undergoing plasticity (Béïque et al., 2011; Lee et al., 2010; Losonczy et al., 2008; Makara et al., 2009).

### Broader applications of zapERtrap

Allowing precise spatial, temporal and gain control, zapERtrap adds critical new capabilities to previously developed “trap and release” strategies for controlling secretory trafficking (Boncompain et al., 2012; Chen et al., 2013; Hangen et al., 2018; Presley et al., 1997; Rivera et al., 2000; Toomre et al., 2000). While we utilized zapERtrap to investigate trafficking from spatially restricted subcellular domains, we expect the approach will have broad appeal because it could be easily implemented for remote control of any secreted substance or for generating user-defined expression patterns of surface proteins with single-cell and subcellular resolution in diverse model systems. For example morphogens, growth factors, endocrine molecules and/or their receptors could be precisely controlled within defined cell populations at specific times. The approach could be used to locally restrict surface expression of receptors with single cell resolution, or to conditionally block surface receptor function by regulated release of inhibitory ligands allowing exquisite spatial and temporal control of cellular signaling events. Thus zapERtrap provides a versatile and robust platform that greatly expands our ability to manipulate protein trafficking and secretion in complex cellular environments.

## Acknowledgements

We would like to thank Mark Dell’Acqua and Chandra Tucker for critical discussions during the preparation of the manuscript. A.M.B. is supported by the National Science Foundation (DGE-1553798) and the Howard Hughes Medical Institute Gilliam Fellowship. A.B.B. was supported by the National Institute of Health (NS092421). Work in the lab of M.J.K. is supported by grants R01NS082271, R01NS10755, R35NS116879 and UF1NS107710 from the National Institute of Neurological Disorders and Stroke.

## Author Contributions

Conceptualization, M.J.K.; Methodology, M.J.K, A.M.B., A.B.B., T.L.S., A.G.; Software, S.L.S. and A.M.B.; Formal Analysis, A.M.B.; Investigation, M.J.K., A.M.B. and A.B.B.; Writing – Original Draft, M.J.K. and A.M.B.; Writing – Review & Editing, M.J.K. and A.M.B.; Funding M.J.K. and A.M.B.; Resources, T.L.S. and A.G.; Supervision, M.J.K.

## STAR Methods

### RESOURCE AVAILABILITY

#### Lead Contact

Further information and requests for resources and reagents should be directed to and will be fulfilled by the Lead Contact, Matthew J. Kennedy.

#### Materials Availability

Plasmids generated in this study will be made available through Addgene.

#### Data and Code Availability

The code generated during this study are available at https://github.com/samanthalschwartz/NeuronAnalysisToolbox/tree/master/zapERtrap.

### EXPERIMENTAL MODEL AND SUBJECT DETAILS

All hippocampal neurons were derived from both male and female neonatal Sprague-Dawley rat pups. Timed pregnant dams (typically embryonic day 16) were obtained from Charles River Laboratories and housed under standard conditions (12 h light/dark cycle, food and water *ad libitum*) until the litter was born. All animal procedures were carried out in accordance with a protocol approved by the University of Colorado Denver Institutional Animal Care and Use Committee.

#### Primary Cell Culture and Transfection

Hippocampi were dissected from P0-1 (postnatal day 0-1) rats and dissociated with 20 units/mL papain (Worthington) in dissociation medium (0.14 M NaCl, 5.4 mM KCl, 0.17 mM Na_2_HPO_4_·7H_2_0, 0.22 mM KH_2_PO3, 9.9 mM HEPES, 44 mM sucrose, 33 Mm glucose, 1.5 mM CaCl_2_, 0.5 mM EDTA, 1 mM NaOH, 0.2 mg/mL cysteine) for 1 h at room temperature and triturated in Minimal Essential Medium (MEM; GIBCO). Cells were plated on poly-D-lysine-coated 18 mm glass coverslips at 25,000 cells/cm^2^ in MEM supplemented with 10% fetal bovine serum (HyClone), 50 units/mL penicillin and 50 μg/mL streptomycin. After 1 d, the medium was replaced with Neurobasal-A (GIBCO) supplemented with B27 (Invitrogen) and GlutaMAX (GIBCO). Neurons were fed with Neurobasal-A containing B27, GlutaMAX, and mitotic inhibitors (uridine + 5-fluoro-2’-deoxyridine [Ur+FdUr]) by replacing half of the medium on day 6 or day 7 and then weekly. Neurons were maintained at 37°C in a humidified incubator at 5% CO_2_. Neurons were transfected between 14 and 18 days in vitro (DIV) with Lipofectamine 2000 (Invitrogen) according to the manufacturer’s recommendations and allowed to express for 24-48 h. For inducible-release experiments, neurons were allowed to express for 24 h and in the presence of 0.5-1 μM zapalog.

#### Cell Line Maintenance and Transfection

COS-7 cells were maintained and propagated in Dulbecco’s Modified Eagle Medium (DMEM; GIBCO) supplemented with 10% fetal bovine serum (HyClone), 50 units/mL penicillin and 50 μg/mL streptomycin at 37°C with 5% CO_2_. COS-7 cells were transfected using Lipofectamine 2000 (Invitrogen) according to the manufacturer’s recommendations and allowed to express for 24 h in the presence of 0.5-1 μM zapalog. COS-7 cell lines were obtained from ATCC, expanded and frozen. Parent cell lines were freshly thawed and validated by cellular morphology, growth characteristics and confirmed to be mycoplasma negative.

### METHOD DETAILS

#### Molecular Cloning

Full sequences of constructs and oligonucleotides used in this study are available upon request. To generate DHFR-tagged cargo molecules, DHFR was inserted following the signal peptide at the N-terminus of FP-NL1 and FP-GluA1 and at the C-terminus of TfR-FP (where FP is EGFP, mCh, halotag, or mNeon) using standard restriction digest cloning or Gibson assembly. We included a thrombin cleavage site between the FP and the open reading frames of the cargo proteins so that any accumulated background surface signal could be eliminated. All constructs were verified by sequencing at the Barbara Davis Sequencing Core at the University of Colorado School of Medicine. The DHHC2-mCh construct was a gift from Dr. Mark Dell’Acqua; The AnkyrinG-mCherry construct was a gift from Dr. Katharine R. Smith; The cDNAs for GluA1 and TfR were gifts from Dr. Michael Ehlers, the NL1 open reading frame was a gift from Dr. Peter Scheiffele’s lab (Addgene clone #15262), the Rab11a reading frame was a gift from Dr. Richard Pagano’s lab (Addgene clone #12674).

#### Imaging

Live cell imaging was performed at 32°C on an Olympus IX71 equipped with a spinning disc scan head (Yokogawa). Excitation illumination was delivered from an acousto-optic tunable filter (AOTF) controlled laser launch (Andor). Images were acquired using a 60x Plan Apochromat 1.4 NA objective and collected on a 1024×1024 pixel Andor iXon EM-CCD camera using Metamorph (Molecular Devices) data acquisition software. For most experiments, a 4.8 μm Z-stack (0.4 μm step-size) was acquired at each time point. For global release experiments, full-field illumination was carried out using a 10-50msec pulse of light at 80% laser power (912 μW measured from the objective) using a 100 mW 405 nm laser. For local release experiments, we focally stimulated the preparation using galvanometric mirrors (FRAPPA, Andor) to steer a diffraction limited 405 nm spot with a 500 μsec dwell time. For all experiments (except for laser intensity titration experiments), local release was triggered with 23.0 μW/μm^2^ 405nm illumination (6% total laser power from a 100mW fiber-coupled laser).

#### Live Cell Surface Labeling and Immunocytochemistry

Neurons were transfected with DHFR-tagged cargoes and FKBP-XFP-KDEL (where XFP is mCh or GFP) using standard lipofectamine2000 (Invitrogen) based transfection protocol according to the manufacturer’s instructions. Zapalog compound (500nM-1μM) was added immediately following transfection and present throughout the experiment. Neurons were imaged in artificial cerebrospinal fluid (ACSF) containing: 130 mM NaCl, 5 mM KCl, 10 mM HEPES, 30 mM glucose, 2 mM CaCl_2_, 1 mM MgCl_2_, 0.5-1 μM zapalog, pH 7.4. Prior to baseline image acquisition, Alexa Fluor 647 anti-GFP (2.67 μg/mL [1:750], Invitrogen) was added to the imaging ACSF for real-time detection of surface accumulation. We typically eliminated minor background signal arising from cargo that escaped the ER by treatment with thrombin (Sigma; cat no. T6884) at 1 unit/mL 10-30 min prior to imaging. We confirmed absence of surface signal arising from recycling or surface receptors that may have escaped the ER by incubating the cells with labeled anti-GFP antibody for a prolonged baseline period (which ranged from 15-20 min) prior to ER release. Cells displaying appreciable surface signal prior to 405nm light exposure were excluded from analysis. For activity dependence experiments tetrodotoxin (TTX; 2 μM final concentration) was added to live cells 30 min prior to imaging and included in the imaging ACSF. Bicuculline (30 μM final concentration) was added to imaging ACSF immediately following ER release. For all experiments we confined our analysis to neurons with a pyramidal-shaped cell body, large apical dendrite and presence of dendritic spines. Primary antibodies used in this study include *α*GFP, Alexa Fluor 555 *α*GFP (Invitrogen; cat no. A31851; 1:500 [4 μg/mL]) and Alexa Fluor 647 anti-GFP (2.67 μg/mL [1:750], Invitrogen). For goat anti-rabbit 568 secondary experiments, both anti-GFP 647 and goat anti-rabbit 568 were used at 1:750. Manual thresholding in ImageJ of 640 image was carried out to create a mask. For some experiments imaging intracellular vesicle formation and movement, we used halo-tagged cargo labeled with JaneliaFluor 646 HaloTag ligand due to its brightness, photostability, and resistance to the low pH in the lumen of trafficking vesicles. JaneliaFluor 646 HaloTag ligand was added to the cell culture media at a final concentration of 100 nM 30 min before live-cell imaging in ACSF. The JaneliaFluor ligand was a gift from Dr. Luke Lavis at the Janelia Farm Research Campus.

### QUANTIFICATION AND STATISTICAL ANALYSIS

All quantification was performed on raw fluorescent images using MATLAB or ImageJ to measure pixel intensities. Background intensity values were estimated either in ImageJ by measuring pixel intensities in image regions with no detectable signal or in MATLAB by interpolating the background intensity within the cell based on the background intensity values outside the cell. Images were expanded in ImageJ for display only. GraphPad Prism 8 was used for performing statistical analyses and plotting data. Heatmaps were generated using MATLAB. For all statistical tests, a p value of less than 0.05 was considered significant. Asterisks denote the following significance levels: *p<0.05, **p<0.01, ***p<0.001, ****p<0.0001.

#### Quantification of Surface Accumulation

Quantification of surface accumulation was carried out using custom analysis software written in MATLAB with the freely available Matlab Toolbox DipImage (Delft University). For each neuron, two masks were generated. The first mask was used to identify the entire cell and used the fluorescent signal of the non-specific cell-fill marker. The second mask identified locations of protein surface accumulation after ER release and was generated using the fluorescent signal of the A647 labeled anti-GFP antibody. For mask generation, images were first filtered using the Laplacian of the Gaussian image transform (implemented by taking only the positive terms from the ‘*dxx*’+ ‘*dyy’* operation in DipImage) followed by a user selected threshold, identified by selecting the maximum intensity within a background region. To quantify the distance of each labeled object along the cell from the cell soma, two approaches were used. Both approaches required a user selected soma region that was drawn as a polygon over the entire somatic body. The first approach quantifies the intensity density of cell surface labeling as a function of distance from the cell soma. A geometric distance transform was applied using the ‘*bwdistgeodesic*’ function in Matlab, with the whole cell mask and identified soma polygon as input. This distance transform was then used to generate a series of masks at variable distances, whose intensity density was used to calculate intensity as a function of distance. Due to non-uniformity of the sample illumination, a custom background was calculated by interpolating the background signal within the cell using the background signal from all regions outside of the cell. Using the Matlab function ‘*regionfill*’ we interpolated a background within the region of the whole cell mask, using background signal. The second approach quantifies the number of aggregated surface receptor puncta as a function of time and distance from the cell soma. To uniquely identify individual puncta, each spatially isolated puncta within the mask was labeled using the DipImage ‘*label*’ function. The distance between each label and the cell soma was calculated by identifying the minimum value after subtracting the geometric distance transform of the soma mask and the geometric distance transform of the individual labeled puncta (found using the Matlab function ‘*bwdistgeodesic*’ as described above), the minimum value representing the minimum path length from the puncta to the soma, along the whole cell mask.

#### Quantification of Surface Accumulation in Dendritic Shafts and Spines

The fraction of cargo-positive spines was calculated within 10 μm regions inside the release zone vs. a region of the same size at a comparable distance from the soma and made from a separate dendrite with similar spine density/morphology.

## Supplemental Movies

**Movie S1. Spatially restricted ER release of TfR-GFP-DHFR from an individual COS-7 cell, related to Figure 1**. Single cell ER release in COS-7 cells expressing TfR-GFP-DHFR (green) and FKBP-mCh-KDEL (channel not displayed) in the presence of 3.33 μg/mL Alexa Fluor 647 anti-GFP (white) before and after focal 405 nm illumination. The photoactivated cell is marked with a purple circle the first frame after photoactivation. Note the accumulation of TfR-GFP-DHFR at the Golgi Apparatus and then the surface in only the targeted cell. The duration of the movie is 50 min with an acquisition rate of 1 frame/2 min. Scale bar, 20 μm.

**Movie S2. Effective retention and light-triggered release from the ER using zapERtrap in a hippocampal neuron, related to Figure 2**. Shown is a hippocampal neuron expressing TfR-GFP-DHFR (green) along with ER-targeted FKBP-mCh-KDEL (red) in the presence of 500nM zapalog before and after a single pulse of full-field 405nm illumination (indicated by the white dot) delivered at time=0 min. Note the rapid light-triggered redistribution of the retained cargo (green) to mobile intracellular dendritic organelles and the stationary GA in the soma. Dimensions of the movie are 82μm x 82μm.

**Movie S3. Kinetics of somatic GA accumulation for different cargoes, related to Figure 2**. Shown are hippocampal neurons expressing FKBP-mCh-KDEL (channel not displayed) and either TfR-GFP-DHFR (left), DHFR-GFP-NL1 (center) or DHFR-mNeon-GluA1 (right) before and after full-field 405 nm illumination (indicated by the white dot) delivered at time=0. The duration of the movies are 67 min with a baseline acquisition rate of 1 frame/min (first 5 frames) and a post-release acquisition rate of 1 frame/2 min. Timestamp displays the time elapsed following full-field 405 nm illumination. Scale bar, 10 μm.

**Movie S4. Surface accumulation of light-released cargoes in neurons** Shown are hippocampal neurons expressing FKBP-mCh-KDEL (channel not displayed) and either DHFR-GFP-NL1 (top) or DHFR-GFP-GluA1 (bottom) before and after full-field 405 nm illumination in the presence of Alexa Fluor647 anti-GFP (the first frame after photoexcitation is denoted by the appearance of the white circle, lower right corner). The duration of the movies are 130 min with a baseline acquisition rate of 1 frame/min and a post-release acquisition rate of 1 frame/2 min. Timestamp displays the time elapsed following full-field 405 nm illumination. Scale bar, 20 μm.

**Movie S5. Segmented surface signal displaying where and when NL1 and GluA1 appear on the cell surface following global release, related to Figure 3**. Surface signal was masked, segmented and displayed only during the first frame of appearance and for 3 subsequent frames (even though the signal was persistent) to better visualize where and when surface signal appeared (rendered in green, generated by a custom MATLAB script, see STAR Methods). The cells were exposed to full field 405nm illumination at t=0min. The cell outlines (purple) were drawn based on a cell fill mask. The first frame after photoexcitation is denoted by the appearance of the white circle, upper right corner). The duration of the movies are 130 min with an acquisition rate of 1 frame/2 min. Scale bar, 20 μm.

**Movie S6. NL1 is inserted into the plasma membrane of the AIS, related to Figure 4**. Shown is a hippocampal neuron expressing DHFR-GFP-NL1 along with unlabeled FKBP-KDEL, ankyrin G-mCh (magenta) and a GFP cell fill (green) before and after full-field 405 nm illumination in the presence of AlexaFluor 647 anti-GFP (surface signal is shown in cyan). The first frame after photoexcitation is denoted by the appearance of the white circle, upper right corner. The duration of the movie is 50 min with a baseline acquisition rate of 1 frame/2 min and a post-release acquisition rate of 1 frame/2.5 min. Timestamp displays the time elapsed following full-field 405 nm illumination at t=0min. Scale bar, 10 μm.

**Movie S7. Following local dendritic release, mobile transport organelles can be transported away from the release zone**. Shown are hippocampal neurons expressing either TfR-GFP-DHFR (top) or DHFR-Halo-NL1 (bottom, labeled with JaneliaFluor 646) along with FKBP-mCh-KDEL (signal not shown). ER-retained cargo was released at time=0 using focally directed 405nm excitation. The timing and location of ER release is shown by purple rectangles. Arrows denote examples of mobile carriers exiting the release zone.

**Movie S8. Surface accumulation of DHFR-GFP-NL1 and DHFR-GFP-GluA1 following local release from the somatic ER, related to Figure 6**. Shown are hippocampal neurons expressing FKBP-mCh-KDEL (channel not displayed) and either DHFR-GFP-GluA1 (top) or DHFR-GFP-NL1 (bottom) along with mCh cell fill (red). ER-retained cargo was released at time=0 min using focally directed 405 nm excitation over the soma. The timing and location of ER release is shown by purple circles before and after focal illumination. The first frame after photoexcitation is denoted by the appearance of the white circle, upper right corner. Blow ups show proximal and distal dendritic regions for each cell (denoted by white boxes). Surface cargoes are visualized with Alexa647 labeled anti-GFP (greyscale). The duration of the movie is 84 min with an acquisition rate of 1 frame/2 min. Timestamps display the time elapsed following focally directed 405 nm illumination. Note the strong surface NL1 accumulation at the putative AIS (arrow) following somatic ER release.

## Figure legends

**Figure S1.**
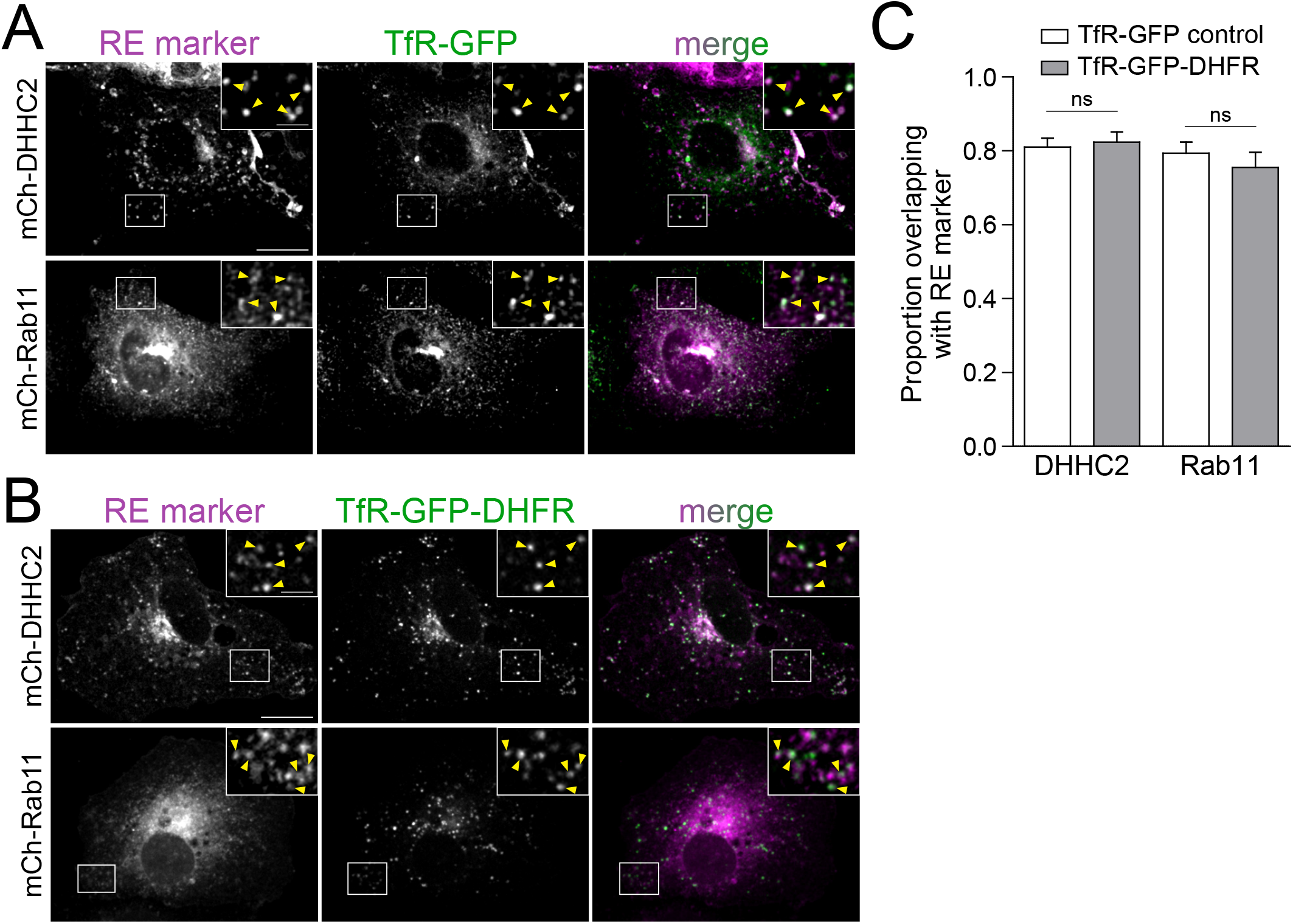
DHFR tag does not alter subcellular localization. **A**. Subcellular localization of TfR-GFP in COS-7 cells relative to the recycling endosome (RE) markers mCh-DHHC2 (*top*) and mCh-Rab11 (*bottom*). Insets show magnified images of the regions marked by the white box. Yellow arrowheads in insets denote colocalized puncta. Scale bar, 20 μm. Inset scale bar, 5 μm. **B**. Subcellular localization of TfR-GFP-DHFR in COS-7 cells relative to the recycling endosome markers mCh-DHHC2 (*top*) and mCh-Rab11 (*bottom*). Insets show magnified images of the regions marked by the white box. Yellow arrowheads in insets denote colocalized puncta. Scale bar, 20 μm. Inset scale bar, 5 μm. **C**. Quantification of the data shown in (A) and (B). Colocalization between TfR-GFP or TfR-GFP-DHFR and the RE markers mCh-DHHC2 and mCh-Rab11 was assessed by calculating the proportion of either TfR-GFP (white bars) or TfR-GFP-DHFR (gray bars) that overlap with the respective RE marker. Data are represented as mean ± SEM. ns=not significant (unpaired t test; p=0.7346 [DHHC2], p=0.4558 [Rab11]; n=10 cells/condition).

**Figure S2.**
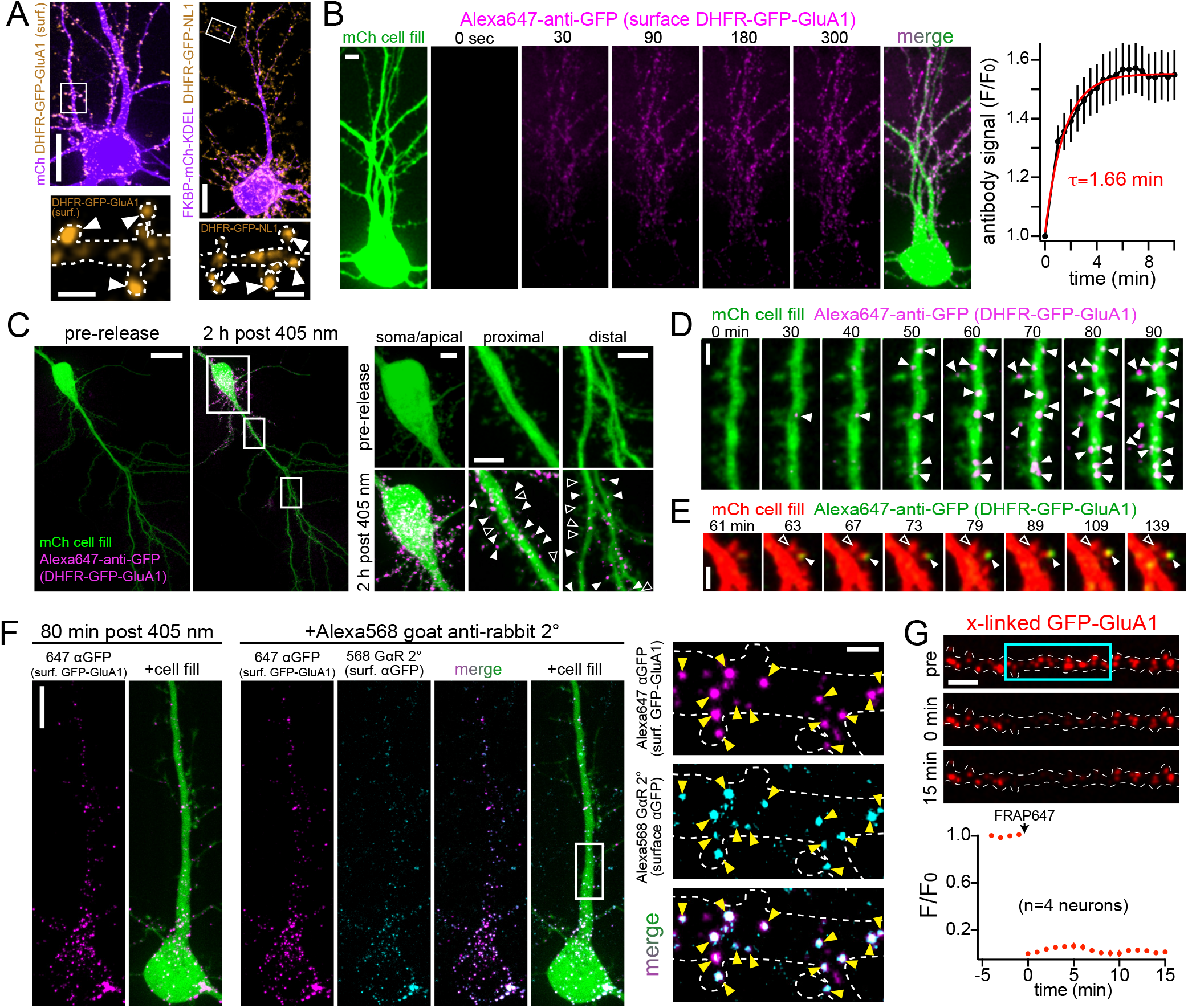
Validation of Antibody crosslinking strategy for visualizing surface accumulation in neurons. **A**. Images of DHFR-GFP-GluA1 (*left*) and DHFR-GFP-NL1 (*right*) under basal conditions, in the absence of zapalog.Scale bars, 20 μm. Inset scale bars, 2 μm. **B**. The binding rate of Alexa Fluor 647-conjugated rabbit anti-GFP (Alexa647-anti-GFP; magenta) was measured in live neurons expressing DHFR-GFP-GluA1. The kinetics of surface binding following addition of the antibody at time 0 are shown in the plot to the right and fitted with a single exponential (tau=1.66 min). Data are represented as mean ± SEM (n=6 neurons). Scale bar, 3 μm. **C**. A neuron expressing DHFR-GFP-GluA1 and a cell fill (green) was imaged before and 2 hours after ER release in the presence of Alexa647-anti-GFP (magenta). Magnified images from different subcellular regions (marked by the white boxes) are shown on the right. Solid arrowheads denote surface-GluA1 containing spines. Open arrowheads mark spines lacking detectable surface DHFR-GFP-GluA1. Scale bar, 10 μm. All inset scale bars, 5 μm. **D**. Image series of DHFR-GFP-GluA1 accumulation on a distal dendritic branch. Note the appearance and stability of new surface clusters (arrowheads) over time. Scale bar, 2 μm. **E**. Image series displaying appearance of DHFR-GFP-GluA1 in a dendritic spine (solid arrowhead) following global ER release. Scale bar, 2 μm. **F**. A neuron expressing DHFR-GFP-GluA1 and a cell fill (green) was imaged 80 min after ER release and following washout of Alexa647-anti-GFP (magenta) that was present in the imaging media (*left*). Following washout of anti-GFP antibody, Alexa568-labeled goat anti-rabbit secondary antibody (cyan) was added and the neuron was imaged 10 min later (90 min after ER release; *right panels*). Scale bar, 10 μm. Magnified images of the highlighted region (white box) are shown to the right. Yellow arrowheads denote colocalized puncta. 86.6 ± 2.83% of anti-GluA1 signal overlapped with anti-rabbit signal. Scale bar, 2 μm. **G**. Shown is anti-GFP signal from surface labeled DHFR-GFP-GluA1. We observed no recovery after photobleaching a section of the dendrite (blue box) confirming effective crosslinking and immobilization. Quantification of Alexa647 signal within the photobleached region is shown below. Data are represented as mean ± SEM (n=4 neurons). Scale bar, 5 μm.

**Figure S3.**
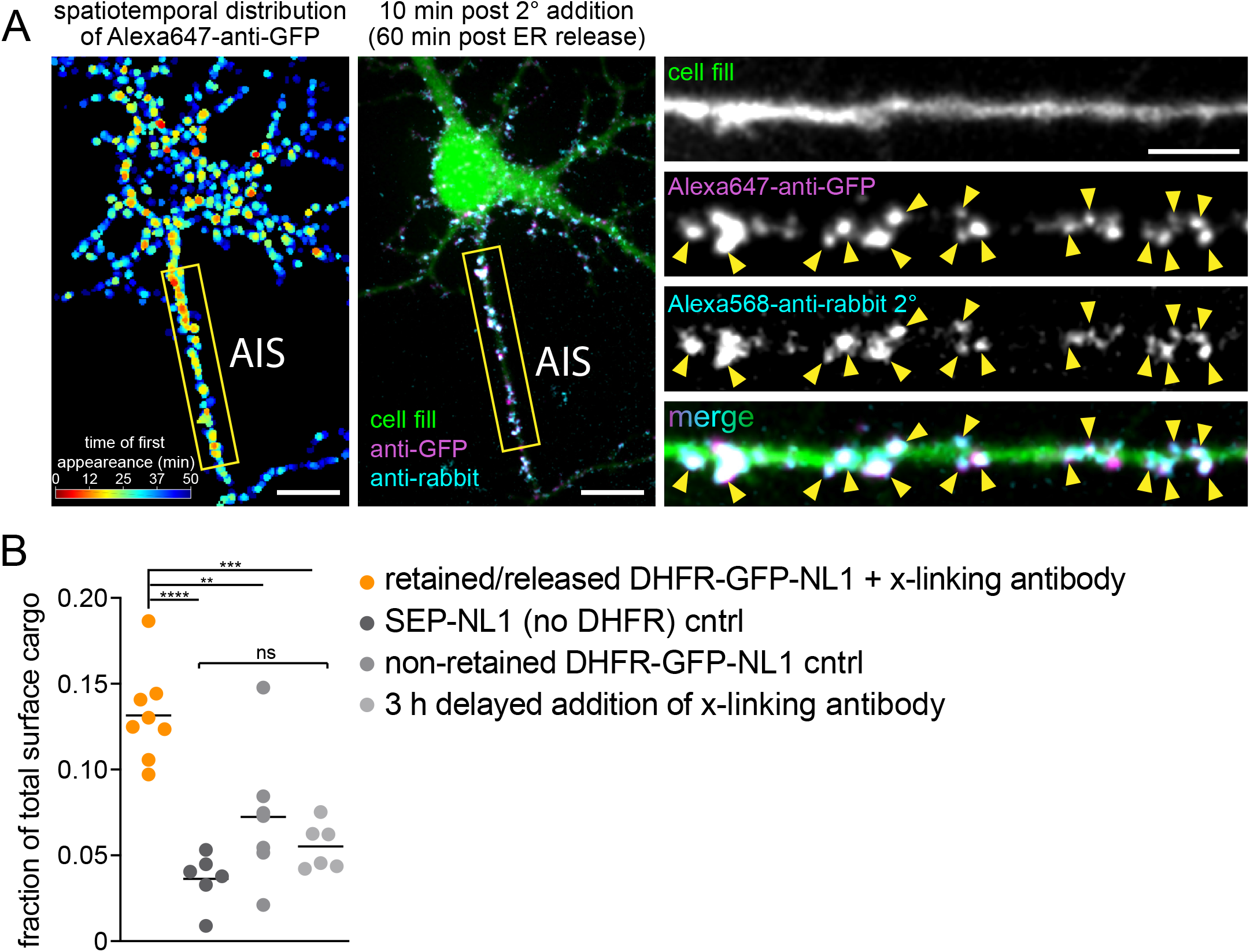
Control experiments for surface NL1 accumulation at the AIS. **A**. NL1 anti-GFP signal is localized at the cell surface. *Left:* Surface accumulation heatmap showing the spatiotemporal distribution of Alexa647-anti-GFP (surface DHFR-GFP-NL1) that was present throughout the ER release experiment and immediately prior to addition of Alexa568-anti-rabbit secondary antibody. *Center:* Confocal image of the same cell 10 min later and following washout of Alexa647 anti-GFP (magenta) and addition of Alexa568 anti-rabbit secondary antibody (cyan). *Right:* Insets of the AIS taken from the yellow box in the image to the left. The robust colocalization of Alexa647-anti-GFP primary and Alexa568-anti-rabbit secondary antibodies (arrowheads) confirm accumulated Alexa647-anti-GFP (DHFR-GFP-NL1) signal is on the cell surface. Scale bars, 10 μm. Inset scale bar, 5 μm. **B**. Comparison of the fraction of total surface cargo at the AIS for retained/released DHFR-GFP-NL1 in the continuous presence of crosslinking antibody (orange) vs. 3 trafficking controls: a SEP-GluA1 (no DHFR) control (dark gray), a non-retained (no zapalog) DHFR-GFP-NL1 control (gray) and DHFR-GFP-NL1 that was released and allowed to traffic for 3 hours prior to addition of crosslinking antibody (light gray). The retained/released DHFR-GFP-NL1 is the same data shown in Fig. 4B with Alexa647-anti-GFP present for the entire duration of the experiment. Data are represented as mean ± SEM. ****p<0.0001, ***p<0.001, **p<0.01, ns=not significant (one-way ANOVA, Tukey’s multiple comparisons test. (n=8 [retained/released DHFR-GFP-NL1]; n=6 [SEP-NL1]; n=7 [non-retained DHFR-GFP-NL1]; n=6 [3 h delayed addition cntrl]; n=number of neurons).

**Figure S4.**
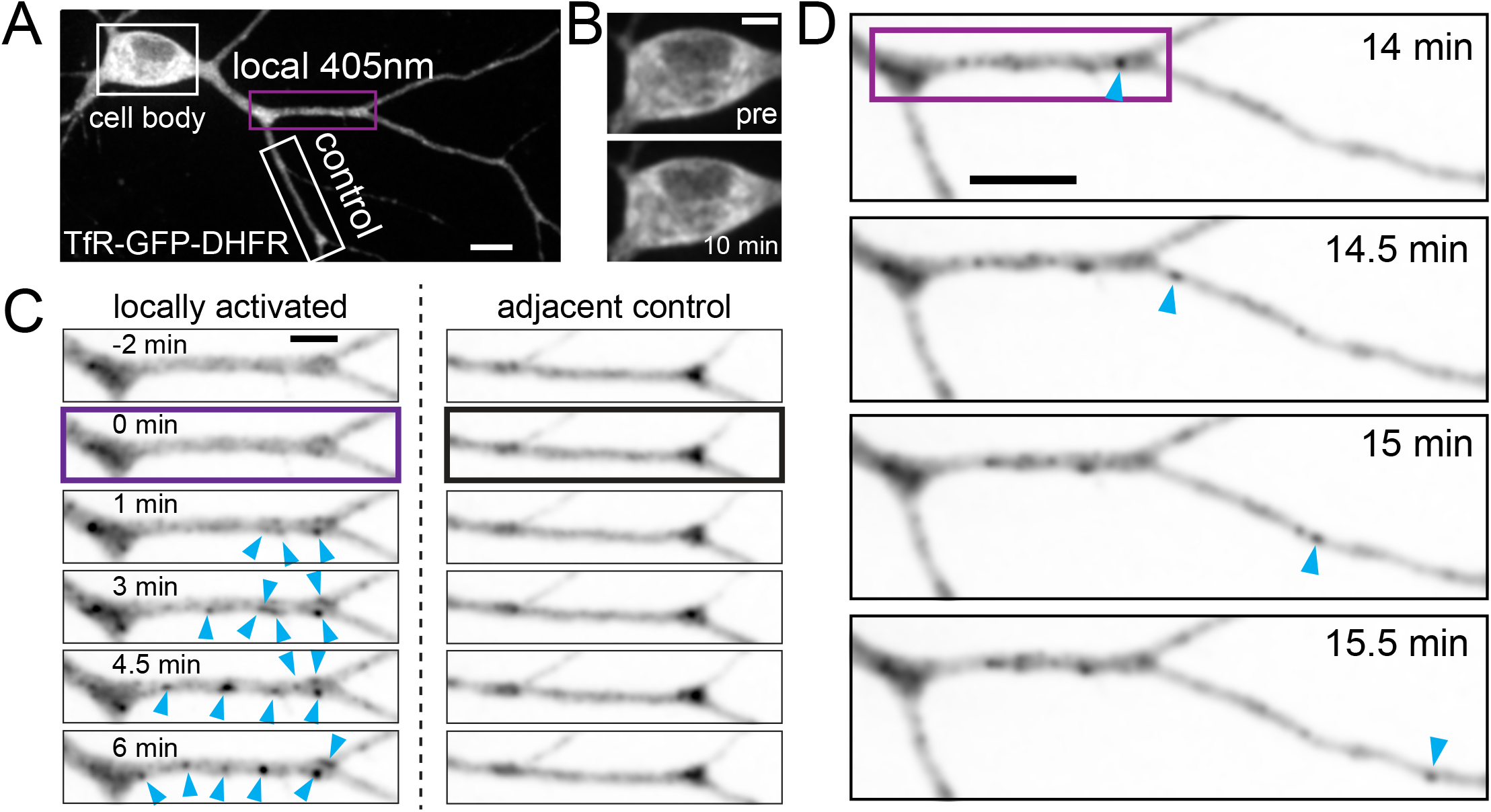
zapERtrap can be used to locally release cargoes from the ER within defined regions of the dendritic arbor. **A**. TfR was released from the dendritic segment under the purple box. The adjacent dendrite serves as a non-photoactivated control.Scale bar, 10 μm. **B**. Somatic TfR-GFP-DHFR signal showed no detectable difference pre and 10 min following local dendritic release. Scale bar, 5 μm. **C**. Time-lapse of intracellular TfR signal following local dendritic release in the targeted branch and an adjacent control branch (white box in [A]). Note the rapid appearance of punctate trafficking organelles (arrowheads) in the released branch but not the control branch. Scale bar, 5 μm. D. Zoomed out images of the dendrite shown in C, starting 14 min after local dendritic photorelease (purple box). An example of a mobile intracellular cargo vesicle (blue arrowhead) exiting the photoactivated region is shown. Scale bar 10μm.

**Figure S5.**
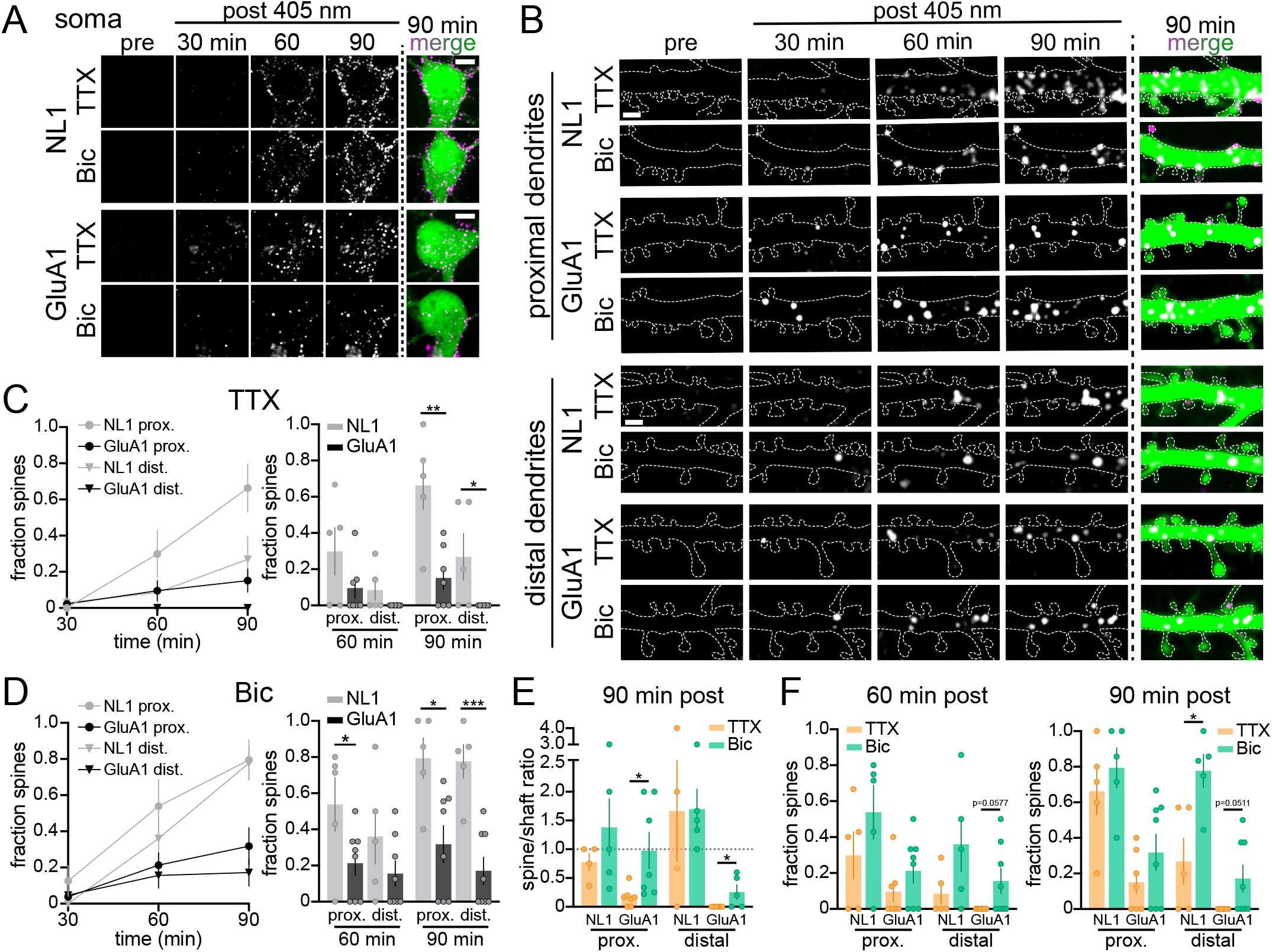
Subcellular distribution of NL1 and GluA1 surface accumulation following somatic ER release. **A**. Image series of DHFR-GFP-NL1 (*top panels*) and DHFR-GFP-GluA1 (*bottom panels*) surface accumulation at the soma before and at 30 min time intervals following somatic ER release in the presence of TTX (2 μM) or bicuculline (bic; 30 μM). Merged images show the Alexa647-anti-GFP signal (magenta) and the cell fill (green). Scale bars, 5 μm. **B**. Image series of DHFR-GFP-NL1 and DHFR-GFP-GluA1 surface accumulation in the proximal (*top panels*) and distal dendrites (*bottom panels*) of the same neurons in (A) before and at 30 min time intervals following somatic ER release in the presence of TTX or bic. Merged images show the Alexa647-anti-GFP signal (magenta) and the cell fill (green). Scale bars, 2 μm. **C**. Time course of the fraction of spines in proximal (circles) and distal (triangles) dendrites that contain surface NL1 (gray) or GluA1 (black) following somatic ER release in the presence of TTX. A comparison of the fraction of NL1- and GluA1-positive spines at 90 min and 120 min is shown on the right. Data are represented as mean ± SEM. **p<0.01, *p<0.05 (unpaired t test; n=5 neurons [NL1], n=7 neurons [GluA1] from at least 2 independent experiments). **D**. Time course of the fraction of spines in proximal (circles) and distal (triangles) dendrites that contain surface NL1 (gray) or GluA1 (black) following somatic ER release in the presence of bic. A comparison of the fraction of NL1- and GluA1-positive spines at 90 min and 120 min is shown on the right. Data are represented as mean ± SEM. ***p<0.001, *p<0.05 (unpaired t test; n=5 neurons [NL1], n=8 neurons [GluA1] from at least 2 independent experiments). **E**. Spine to shaft ratios of surface NL1 or GluA1 signal in proximal and distal dendrites 90 min following somatic ER release in the presence of TTX (orange) or bic (green). Note the split y-axis. Data are represented as mean ± SEM. *p<0.05 (unpaired t test; n=4-5 neurons [NL1], n=5-7 neurons [GluA1] from at least 2 independent experiments). **F**. Comparison of the fraction of cargo-positive spines for NL1 vs. GluA1 at 60 min (*left*) and 90 min (*right*) following somatic ER release in the presence of TTX (orange) or bic (green). Data are represented as mean ± SEM. *p<0.05 (unpaired t test; n=5 neurons [NL1], n=7-8 neurons [GluA1] from at least 2 independent experiments).

